# *Antennapedia* and *optix* regulate metallic silver wing scale development and cell shape in *Bicyclus anynana* butterflies

**DOI:** 10.1101/2021.10.19.464939

**Authors:** Anupama Prakash, Cédric Finet, Tirtha Das Banerjee, Vinodkumar Saranathan, Antónia Monteiro

## Abstract

Butterfly wing scale cells can develop very intricate cuticular nanostructures that interact with light to produce structural colors including silvery hues, but the genetic basis of such nanostructures is mostly unexplored. Here, we first identified that *optix* is necessary for metallic scale development in the butterfly *Bicyclus anynana*. We then characterized different subtypes of wildtype metallic silver scales and addressed the function of five genes – *apterous A, Ultrabithorax, doublesex, Antennapedia*, and *optix –* in the differentiation of silver scales, at a single-cell resolution, by leveraging crispants that exhibited either ectopic gains or losses of silver scales. Wildtype silver scales were generally rounded and had low amounts of pigmentation, exhibiting a common ultrastructural modification for metallic broadband reflectance, i.e., an undulatory air layer enclosed by an upper and lower lamina. Our results indicated that the varying air layer thickness was the important parameter of the bilaminate scale for producing a broadband reflectance across visible wavelengths. While a single lamina of the appropriate thickness could also produce broadband colors, the bilaminate structure is advantageous as it increases the overall reflectivity. Crispant brown scales differed from wildtype silver scales via the loss of the continuous upper lamina, increased lower lamina thickness, and increased pigmentation. The reverse was seen when brown scales became silver. We identified *Antennapedia* and *optix* as high-level regulators in the network differentiating different silver scale subtypes and determining overall cell shape in both sexes. In addition, *Antp* exhibits a novel, post-embryonic role in the determination of ridge and crossrib orientation.

## Introduction

Silver or gold colors in insect cuticles (1–3), fish scales (4), and the eyes of cephalopods (1, 4–6) are all examples of naturally occurring broadband structural coloration. These colors serve multiple ecological functions including in vision, as inter-or intra-specific signals, or in thermoregulation (5, 7–12) and arise from the interaction of light with specific classes of broadband reflectors found in the animal integument (13–15). Broadband metallic reflectors that are highly reflective across a broad range of wavelengths, usually involve thin film or multi-layer interference made of alternating materials with different refractive indices and varying thicknesses (1, 16). Examples of such multi-layer broadband reflectors are found in the exocuticle of gold beetles (16, 17), the endocuticle of some butterfly pupal cases (3, 18, 19), and in fish scales (6, 20). Due to the small difference in refractive indices of biological materials, broadband multilayer reflectors with high reflectivity usually require a minimum of 10-20 alternating layers of high- and low-refractive index materials, leading to very thick (tens of microns) multilayer reflectors (19).

In contrast, broadband metallic reflectors found in lepidopteran wing scales are anatomically constrained by the thickness of the scales. These metallic reflectors are ultra-thin, with an overall thickness of a few microns (10, 21–24). An essential modification of the basic scale Bauplan to produce such broadband reflectors is the consistent presence of a contiguous upper lamina that closes the normally “open” windows seen in a typical scale (10, 21, 24, 25). This creates an undulatory air layer sandwiched by the lower and upper laminas whose thicknesses also vary spatially. Broadband reflectance is achieved by additive color mixing that occurs due to local spatial variation or disorder in the scale ultrastructure (10, 21, 23, 24, 26). The broadband metallic reflectors seen in fossil moths (23, 27), in extant moths (22), and in springtails (28), however, utilize thin film interference from a single chitin layer, resulting from fused scales. Such a single thin film is also present as the lower lamina of archetypal butterfly scales which are often also tuned to produce broadband colors (29, 30). Therefore, the necessity and advantage(s) of a bilaminate system in broadband scale color generation – a trait that has repeatedly evolved across butterflies – is enigmatic.

The production of metallic broadband reflectors may also depend on the presence or absence of pigments embedded in the scale cuticle. For instance, the brown ground scales in *Hypolimnas salmacis*, the yellow scales in *Heliconius* butterflies, and the “black” scales in the *yellow* mutant of *Bicyclus anynana* butterflies have a closed upper lamina (31–33), but these scales do not exhibit broadband reflectance either due to the high concentration of pigments, or incorrect thicknesses of both the laminas and air layer in these scales. These examples suggest that the regulatory networks that create a metallic scale type must involve regulation of scale ultrastructure dimensions perhaps along with pigmentation. While the optical origin of broadband silvery colors from thin-film and chirped multi-layer reflectors in the arthropod integument is well understood (3, 10, 19, 21, 23, 26), the necessity of a bilaminate air-filled nanostructure, the relative contributions of air gap layer vs upper and lower laminas, and the genetic circuits that create these broadband metallic reflectors in butterfly wing scales remain unclear, despite a number of studies on silvery butterfly scales (10, 21, 24, 26)

We have begun to explore the genetic basis of silver scale development with a series of CRISPR experiments performed in the nymphalid butterfly *B. anynana*. This species exhibits different subtypes of broadband reflecting metallic scales on both fore- and hindwings (34). These scales are usually associated with the sex pheromone producing regions in males (the androconia), except for the coupling scales near the base of the wings that are present in both sexes. Previously, the knockout of four genes including *apterous A* (*apA*) (35), *Ultrabithorax* (*Ubx*) (36), *doublesex* (*dsx*) (34), and *Antennapedia* (*Antp*) (36) led to phenotypic effects on metallic scale development in *B. anynana* (Fig 1A). In addition, we speculated that *optix* might be a good candidate regulating metallic coupling scale development in *B. anynana* given that the expression and function of *optix* in the development of coupling scales has previously been documented in other butterflies (37, 38). All modified scales provided an opportunity to investigate the structural basis of the different metallic colors and understand the roles of these genes in creating broadband reflectors in lepidopterans.

**Figure 1:**
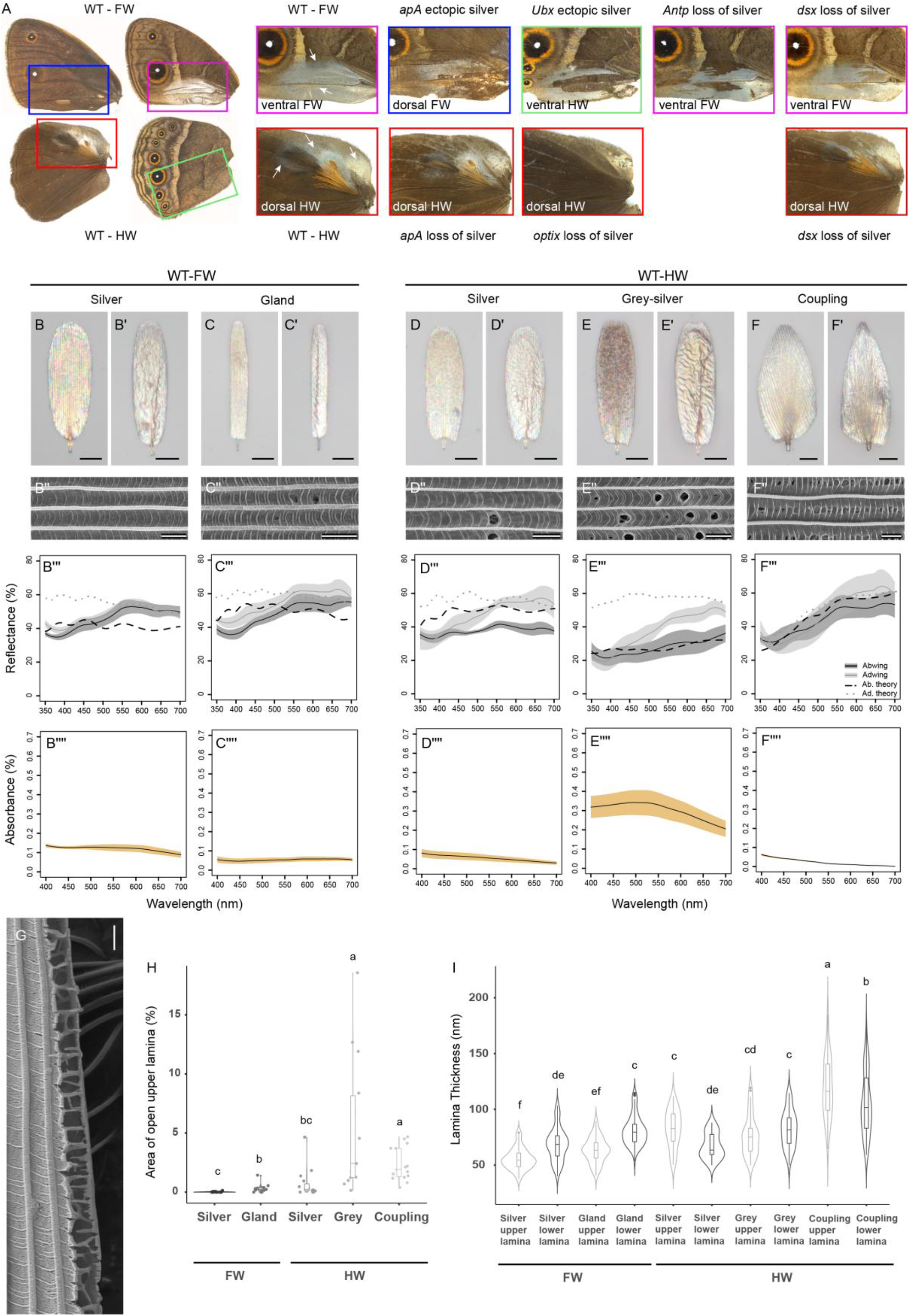
*B. anynana* crispants sampled in this study and broadband reflectors in wildtype *B. anynana*. (A) WT dorsal and ventral wing surfaces with magnified views of the silver scale regions alongside ectopic silver scales or silver to brown transformed scales in crispants of the five genes. The colored box outlines and the text label in each inset indicate the position on the different wing surfaces of the WT and crispant scales. White arrows in the WT panels indicate the five different silver scale types characterized. Optical microscopy images of the abwing (upperside) (B-F) and adwing (underside) (B′-F′) surfaces of single scales, SEM images of the abwing surface (B′′-F′′), measured and modeled reflectance (B′′′-F′′′) and absorbance spectra (B′′′′-F′′′′) of the WT forewing silver and gland scales and the hindwing silver, grey-silver and coupling scales respectively. (G) A longitudinal cross-section of a forewing silver scale showing the upper and lower laminas enclosing an air layer. (H) Area of the open upper lamina of the different silver scales on the forewings and hindwings. Boxplots show the median, inner and outer quartiles and whiskers up to 1.5 times the inter-quartile range. (I) Violin plots of the upper and lower lamina thicknesses of the different forewing and hindwing silver scales. Means sharing the same letter are not significantly different (Tukey-adjusted comparisons). Scale bars: B,B′-F,F′ are 20 µm and B′′-F′′, G are 2 µm.

Among the genes investigated, *apA* and *Ubx* are well known homeotic genes that specify wing or wing-surface identities in butterflies (35, 36, 39, 40). In *B. anynana, apA* is expressed in dorsal wing surfaces and *Ubx* across the entire hindwing, and specify the respective identities of those tissues (35, 36). In contrast, *Antp* and *dsxM* show strong punctate expression within individual scale cells in a trait-specific manner including in eyespot centers, pheromone glands, and the silver scale regions (34, 36) hinting at a functional role of these two genes in differentiating specific scale types. Here, we asked whether the investigated genes modified different structural and pigmentary aspects and components of the scale to create a silver color, and vice versa, when silver scales became brown. We examined whether transformations of different structures and levels of pigmentation within a scale, with the manipulation of each gene in isolation, is complete or gradual. We hypothesized that the two homeotic genes would produce more complete and extreme transformations of the scales, making them resemble a wildtype silver scale (or a brown scale) more closely while the trait-specific genes would have more restrictive transformations, affecting either aspects of color or morphology only.

To answer these questions, we first examined the potential role of *optix* in the development of silver-colored coupling scales in *B. anynana* using immunostainings and CRISPR and identified that *optix* is necessary for coupling scale development. We then used light, conventional and focused ion beam (FIB) scanning electron microscopy (SEM), UV-VIS-NIR microspectrophotometry, and systematic optical modeling, to understand different aspects of the wildtype (WT) and transformed scales for all five genes. We show that the bilaminate structure of metallic scales increases broadband reflectivity in contrast to a single thin-film and that the variable air layer thickness in silver scales is the most important determinant of the broadband nature of the reflectance. Transformation of silver to brown scales is accompanied by the loss of the contiguous upper lamina, increase in lower lamina thickness and gain in pigmentation. The opposite occurs when brown scales become silver scales. In addition, we identify *Antp* and *optix* as high-level regulators of the different silver scale subtypes, determining both scale shape and scale color via modulation of ultrastructure dimensions.

## Results

### *optix* is required for the development of silver colored, trowel-shaped coupling scales in *Bicyclus anynana*

We first investigated the role of *optix* in coupling scale development using immunostainings and CRISPR-Cas9 gene knockouts. Optix protein was highly expressed in punctate nuclei of the future coupling scales on the developing pupal dorsal hindwings at all three time points investigated (Supp Fig 1A-D). Mosaic crispants that resulted from embryonic injections of two guide RNAs against *optix* revealed a silver to brown color transformation of the coupling scales along with a change in shape from typical trowel-shaped scales to scales with rounded or slightly scalloped distal edges (Supp Fig S1E – J). These results indicate that, like *Antp* and *dsx* (34, 36), *optix* is expressed in a trait-specific manner and is required for regulating the color and shape of the silver coupling scales in *B. anynana*.

### Ultrastructural and optical origin of broadband silver reflections in *B. anynana* butterflies

We then investigated how the silver color is produced in five subtypes of silver scales present on the wings of *B. anynana* using light reflection and absorbance measures, as well as SEM (Fig. 1B-G). On the ventral forewing, we focused on the silver and the gland scales on the androconial patch of males (Fig 1A, WT-FW arrows). On the dorsal hindwing, we sampled the silver and grey-silver scales near the androconial patches of males. Finally, we sampled coupling scales near the base of the dorsal hindwing in females (Fig 1A, WT-HW arrows). All silver scales generally feature rounded edges. The coupling scales, however, have a characteristic trowel-head shape (Fig 1F, F′), and their ridges are oriented at an angle to the proximo-distal axis of the scale as compared to the parallel arrangement of ridges in most other scale types. The abwing and adwing surfaces of the scales exhibited an oil-slick like multi-hued appearance, typical of a thin film with varying thickness (Fig 1B-F). The reflectance spectra of all silver scales exhibited broadband (relatively flat or uniform) reflectance, with higher intensities from the adwing surface (Fig 1B′′′-F′′′). Pigmentation levels were low in all scales, as measured from low light absorbance (Fig 1B′′′′-F′′′′), with the exception of the grey-silver scales which absorbed more light, indicating higher levels of pigmentation. All silver scales exhibited a closed upper lamina (Fig 1B′′-F′′), creating a bilaminate structure enclosing an air layer (Fig 1G). The different silver scale subtypes exhibited varying levels of perforations in the total area of upper lamina (Fig 1H, Supplementary Table S5 – source data): from ∼0% (fore- and hindwing silver scales) and 2.35% (hindwing coupling scales), to 5.4% open (hindwing grey-silver scales) (Supplementary Tables S1,2). This correlation between the integrity of the upper lamina coverage to the production of broadband reflectance affirms the important role attributed to the upper lamina and/or the enclosed air layer.

In order to theoretically model these broadband metallic colors, we measured the air gap, lower and upper lamina thicknesses from FIB-SEM cross-sections of the different silver scale types (Fig 1G, I, Supplementary Table S5 – source data). The mean upper and lower lamina thicknesses of the different metallic scale types ranged from approximately 55-120 nm, with the coupling scales having the thickest lower and upper laminas (Fig 1I). We modeled the theoretical adwing and abwing spectra, the latter by incorporating the measured absorptivity curves (Fig 1B′′′′-F′′′′), which produced broadband reflectance with a series of very shallow peaks, whose positions are largely consistent with those in the corresponding measured spectra (Fig 1B′′′-F′′′, Supp Fig S2). A lower lamina thin film of an appropriate thickness could, alone, produce a broadband silver color (Supp Fig S3A). However, our results indicated that the presence of the additional upper lamina enclosing an air gap with varying thickness increased the mean broadband reflectivity by around 11% (Supp Fig S3A). Furthermore, our hierarchical modeling (Supp Figs S3 and S4) suggests that the varying air gap layer is the most important component of the bilaminate 3-layer nanostructure responsible for a broadband silvery reflectance. For instance, the simulated patches with distinct colors (e.g., purple-pink and blue-green; Supp Fig S3C) not only correspond well to those seen in between ridges in high magnification light micrographs (Fig 1B,B′-F,F′), but also combine to make a broadband color in the far-field in a Pointillist fashion (24). However, changing the upper and/or lower lamina thicknesses with a constant air gap thickness cannot reproduce the relatively flat broadband reflectance, even when accounting for the increased lamina thicknesses (i.e., greater variability) at or near the ridges (cf. Supp Figs S3A and S3D).

Interestingly, our systematic parametric modeling (Supp Fig S4) revealed that the thicknesses of both upper and lower laminas for each of the silver scale subtypes, except for the coupling scales, are more or less optimized to produce the maximum broadband reflectivity. By contrast, the air gap layer is sufficiently thick (> 750 nm) to ensure the broadband nature of the reflectivity. A thickness of less than 500 nm would produce strong peak(s) with chromatic effects destroying the silvery appearance (Supp Fig S4). The modeling results also predicted lower reflectivity at short wavelengths below 450 nm for the coupling scales (Supp Fig S4N), like their measured spectra (Fig 1F′′′). Given that the coupling scales are among the least pigmented of the silver scales (Fig. 1F′′′′), our results indicate that this lower reflectivity is a previously undocumented structural rather than a pigmentary effect. The lower reflectivity is caused by a decrease in the thickness of the lower lamina from base to tip (Supp Fig S5) that results in a concomitant change in the color of the scale from bronze/brown in the base/center to silver at the tips. The increased thickness near the base of the scale suppresses shorter wavelengths of light and leads to a greyish darker color (Fig 1F′), while optimum thickness towards the tip reinforces broadband silver reflectance.

### *Antp* regulates silver scale shape and color but *dsxM* only regulates scale color

Both *Antp* and *dsx* male crispants exhibited a silver to brown color transformation on the forewing but had different effects on scale shape (Fig 2A-C). The rounded edges of the distal end of the silver scales of *Antp* crispants changed into serrated finger-like projections, resembling other brown scales on the same wing surface (Fig 2D, Supp Fig S6A). This was also true in *Antp* crispant females, where the normal rounded brown scales on the posterior region of the ventral forewings became brown scales with dentate margins in the crispants (Supp Fig S6B). Furthermore, some of the *Antp* crispant brown scales exhibited unusual ridge and crossrib orientations near the distal tips of the scales, often lying perpendicular to the normal parallel arrangement of ridges (Fig 2F). In contrast, the loss of *dsxM* expression in the same region in males produced brown scales with a rounded distal edge (Fig 2E). Loss of *dsxF* in females had no effects on scale color or shape in the homologous regions. On the hindwing silver areas, we noticed similar small clonal patches of brown scales in *Antp* male crispants (Fig 2G). Such clonal patches were also visible on female hindwings of *Antp* crispants in homologous regions of the wing, though not as contrasting (Supp Fig 6D). The brown scales in these clonal patches had a dentate morphology like other dorsal hindwing scales, in contrast to the rounded scales seen in the hindwing silver scale region in males or the homologous regions in females (Supp Fig S6C, D).

**Figure 2:**
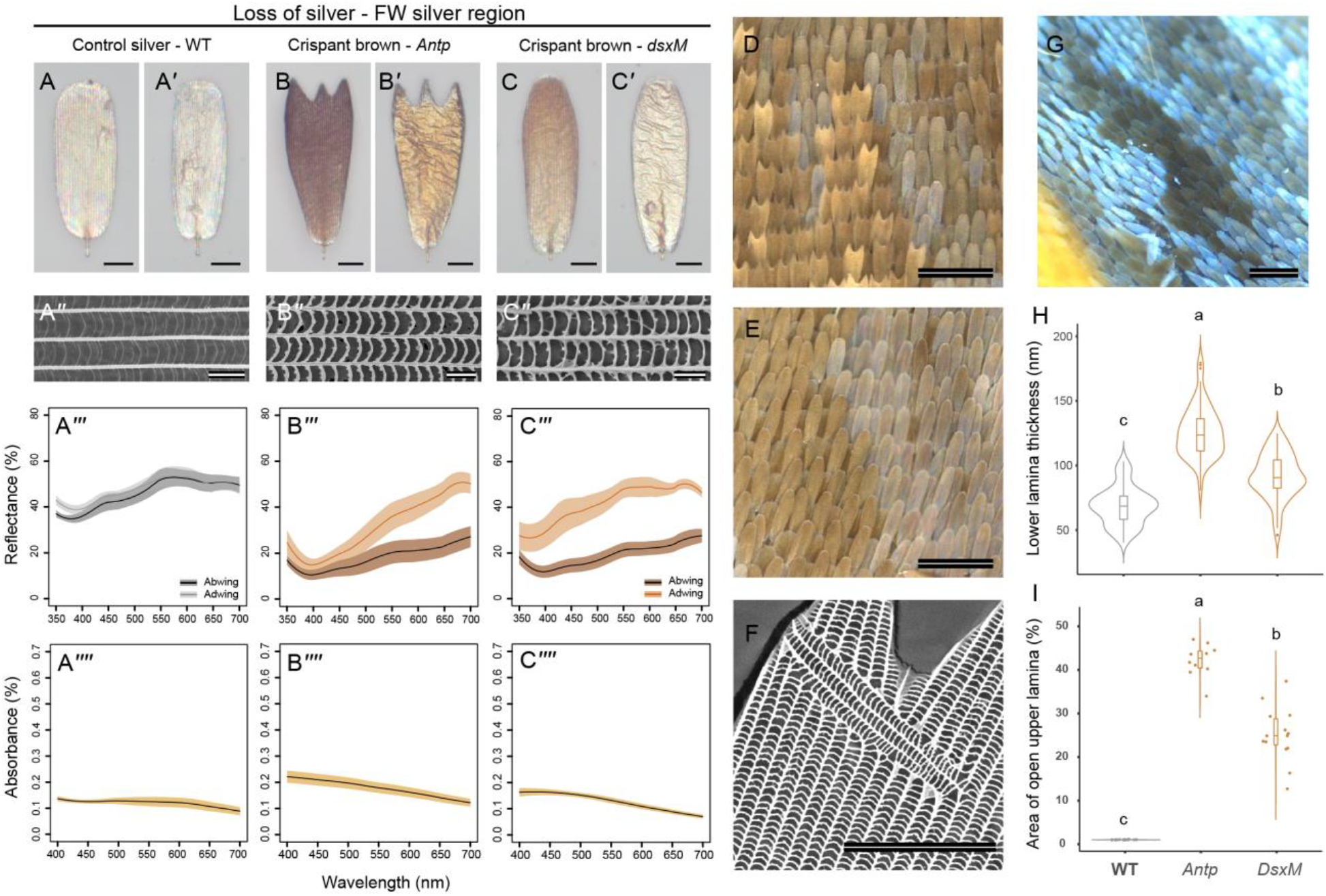
Characterization of the ultrastructure and pigmentation of the silver to brown scales in *Antp* and *dsxM B. anynana* crispants. Optical microscopy images of the abwing (A-C) and adwing (A′-C′) surfaces, SEM images of the abwing surface (A′′-C′′), reflectance (A′′′-C′′′) and absorbance spectra(A′′′′-C′′′′) of the WT forewing control silver scales and the mutant brown scales of *Antp* and *dsxM* crispants respectively. (D) Mosaic scale phenotype in *Antp* crispant forewing illustrating the mutant brown scales with dentate sculpting in the distal tips. (E) Mosaic scale phenotype in *dsxM* crispant forewing illustrating the mutant brown scales with rounded distal tips. (F) SEM image of an *Antp* crispant forewing brown scale showing a nearly orthogonal ridge and crossrib orientation among the normal ridges and crossribs. These variations always occur at the distal tips of the crispant scales (N=4 scales from one individual). (G) Mosaic scale phenotype in male *Antp* crispant hindwing illustrating the mutant brown scales with dentate sculpting in the distal tips. (H) Violin plots of the lower lamina thicknesses of the control silver scales and the crispant brown scales of *Antp* and *dsxM* crispants (I) Total area of open upper lamina of the different control and mutant scales. Boxplots show the median, inner and outer quartiles and whiskers up to 1.5 times the inter-quartile range. Means sharing a letter are not significantly different (Tukey-adjusted comparisons). Scale bars: A,A′-C,C′ and F are 20 µm, A′′-C′′ are 2 µm, D, E, G are 200 µm.

Ultrastructure and pigmentation modifications accompanied the loss of silver reflectance in male scales of both crispant types. The brown crispant scales had lower reflectivity compared to the wildtype control scales (Fig 2A′′′-C′′′) and this was accompanied by an increase in absorbance due to greater pigmentation (Fig 2A′′′′-C′′′′). The crispant brown scales also lost their continuous upper lamina which now exhibited perforated windows (Fig 2A′′-C′′, I, Supplementary Tables S1,2). Other ultrastructural modifications included an increase in lower lamina thickness (Fig 2H) (Posthoc tests from LME, adjusted P-values <0.001; Supplementary Tables S3,4). *Antp* brown scales had a greater lower lamina thickness (125.82 nm ± 19.7 nm) compared to the *dsxM* brown scales (91.9 nm ± 17.6nm) and both were significantly different from wildtype silver lower lamina thickness (68.8 ± 14.75). The increase in pigmentation, loss of the upper lamina and increased lower lamina thickness led to the loss of broadband metallic silver reflectance and created a brown color due to the combination of pigments and the now dominant lower lamina reflectance (Fig 2A′′′-C′′′).

These results suggest that *Antp* is an important high-level regulator in the silver scale differentiation program and is necessary to determine correct silver scale color and morphology in males and scale shape in females. *Antp* likely acts upstream of, or in parallel with *dsxM*, which appears to have similar effects on color and upper lamina morphology but does not impact scale shape. In addition, *Antp* crispant scales showed larger changes in lower lamina thickness and area of open upper lamina compared to *dsxM* crispant scales (Fig 2H, I). To investigate the extent to which Antp expression maps to silver scales, we examined the expression of Antp in 24-hour male and female pupal wings. Antp protein was visible in punctate nuclei in the forewing (Supp Fig S7A) and hindwing silver scale regions in males, especially around the future hindwing gland (Supp Fig S7C), and also in female wings at homologous locations (Supp Fig S7B, D). Antp protein expression was lower in the grey-silver scales on male hindwings (Supp Fig S7C). This pattern on male hindwings was especially clear in a 48-hour pupal hindwing (Supp Fig S8). These data indicate that Antp protein maps precisely to punctate nuclei in the area where silver scales develop on both wings, as expected of a transcription factor.

### *apA, dsxM*, and *optix* promote silver coloration via the gain of an upper lamina, decrease in lower lamina thickness and loss of pigmentation

On the hindwings, crispants of *apA, dsx*, and *optix* showed loss of metallic silver scales and transformation of these scales into brown scales. For *apA* these are the exact opposite phenotypes from those observed on the forewing, where *apA* transformed brown scales into ectopic silver scales. In *apA* and *dsxM* crispants, hindwing silver and grey-silver scales were transformed into brown scales (Fig 3A-F). Brown crispant scales of *apA* individuals exhibited a dentate morphology while those in *dsxM* individuals exhibited some variation in morphology, from rounded scales in the silver region to sometimes dentate scales in the grey-silver region. Crispants of *optix* affected both the shape and the color of coupling scales (Fig 3G-H, Supp Fig 1H – J). Reflectance spectra reflected the change in coloration (Fig 3A′′′ – H′′′). In all instances of transformation, the adwing sides of the crispant scales were thin films reflecting a deep bronze-golden color instead of the silver to gold reflectance of the wildtype control scales (Fig 3A′-H′). In addition, in line with the silver to brown transformation, there was an increase in pigmentation in most scales (Fig 3A′′′′-C′′′′, G′′′′-H′′′′), except in the grey-silver region, where crispant brown scales, in both *apA* and *dsxM* crispants, had decreased pigmentation as compared to the controls (Fig 3D′′′′-F′′′′), despite control scales displaying a lot of variation in pigmentation levels (Fig 3D′′′′).

**Figure 3:**
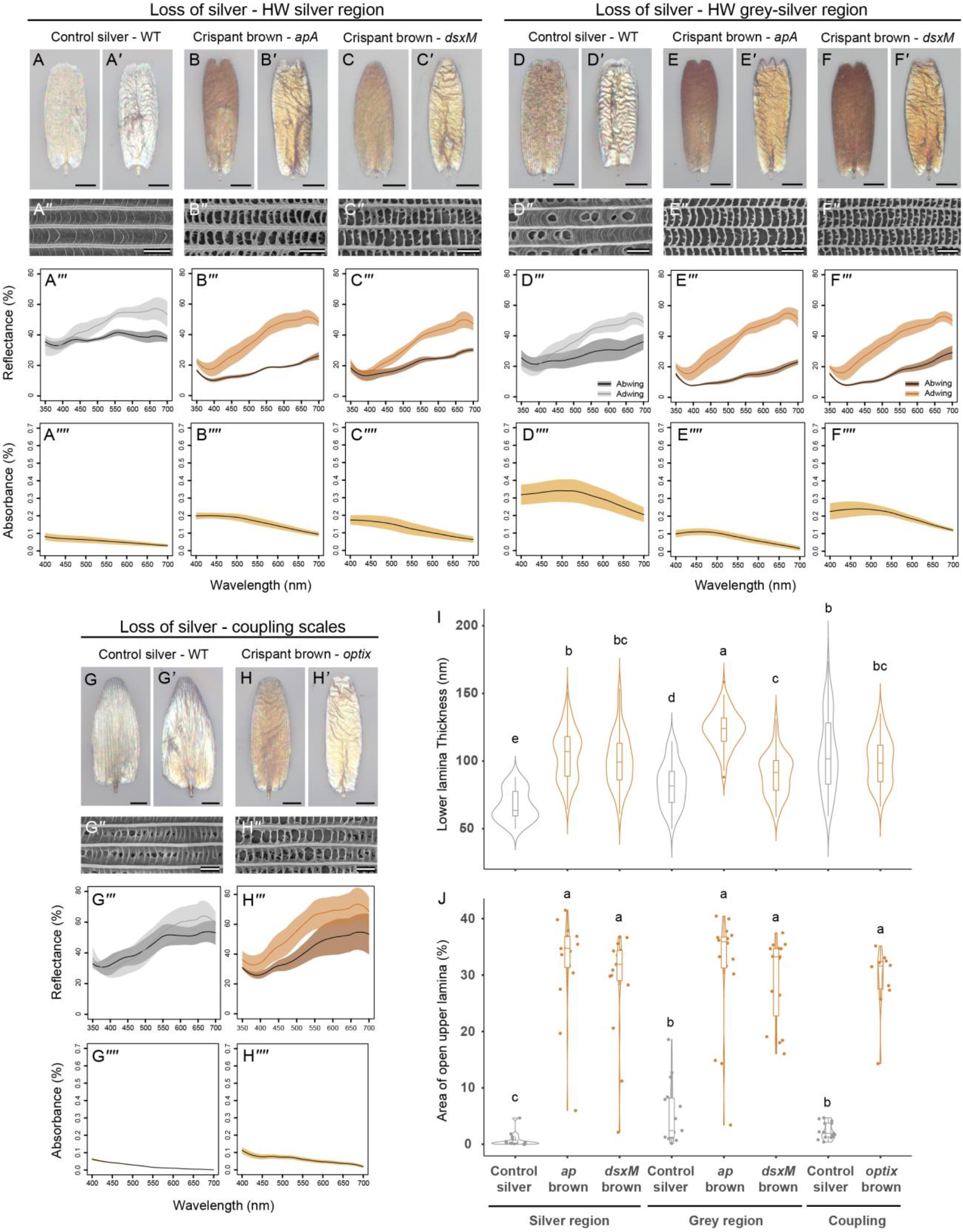
Characterization of the ultrastructure and pigmentation of transformed silver to brown scales in *apA, dsxM* and *optix B. anynana* crispants. Optical microscopy images of the abwing (A-H) and adwing (A’-H’) surfaces, SEM images of the abwing surface (A′′-H′′), reflectance (A′′′-H′′′) and absorbance spectra(A′′′′-H′′′′) of the control WT silver scales and transformed crispant brown scales in the hindwing silver region (A-C), the hindwing grey-silver region (D-F) and the hindwing coupling scales (G-H). The control scales are WT silver scales from homologous regions on the wing. (I) Violin plots of the lower lamina thicknesses of control WT silver scales and the crispant brown scales of *apA, dsxM* and *optix* crispants (G) Total area of open upper lamina of the different control and crispant brown scales. Boxplots show the median, inner and outer quartiles and whiskers up to 1.5 times the inter-quartile range. Means sharing a letter are not significantly different (Tukey-adjusted comparisons). Scale bars: A,A′-H,H′ are 20 µm and A′′-H′′ are 2 µm.

Crispant brown scales on the hindwing, produced by all three genes, also exhibited a loss of the upper lamina (Fig 3A′′-H′′, J, Supplementary Tables S1,2). This was not a clean morphological change, with many scales exhibiting a gradualism in transformation, often having remnants of the lamina attached to the crossribs and ridges (Fig 3A′′-H′′, J). The lower lamina thickness of these scales also increased compared to wildtype silver controls (Posthoc tests from LME, adjusted P-values <0.001; Supplementary Tables S3,4). Both *apA* and *dsxM* hindwing brown scales in the silver and grey-silver regions had significantly thicker lower laminas than control scales, while there was no change of lamina thickness in *optix* crispant brown scales (Fig 3I, Supplementary Tables S3,4).

### Ectopic silver scales gain an upper lamina, have a thinner lower lamina and lose pigmentation

We next investigated the ectopic forewing silver scales produced in the *apA* and *Ubx* crispants by the transformation of non-reflective brown scales (Fig 1A). We characterized both ectopic silver and gland scale subtypes in *apA* crispants but only the ectopic silver scale subtype in the single *Ubx* crispant due to size limitation of the mutant clone in that individual. Ectopic silver scales in both *apA* and *Ubx* crispants looked like wildtype silver scales, including changes in cell shape (Fig 4). The *Ubx* crispant scales exhibited an extensive transformation, from scalloped brown scales to rounded silvery scales (Fig 4D, E), while the *apA* crispant scales changed primarily in color and less extensively in shape (Fig 4A-C). The abwing and adwing surfaces of the ectopic scales were metallic silvery thin films (Fig 4B, C, E). These scales were dramatically different from the control brown scales from the same crispant regions which were brown in color from the abwing side (Fig 4A, D) and had a bronze-golden adwing thin film (Fig 4A′, D′). Reflectance spectra corresponded with the optical images, with the silver scales reflecting broadly (Fig 4A′′′-E′′′). Ectopic metallic scales of both *apA* and *Ubx* crispants also lost pigmentation as compared to the control brown scales, though there was variation between the different scale subtypes (Fig 4A′′′′-E′′′′).

**Figure 4:**
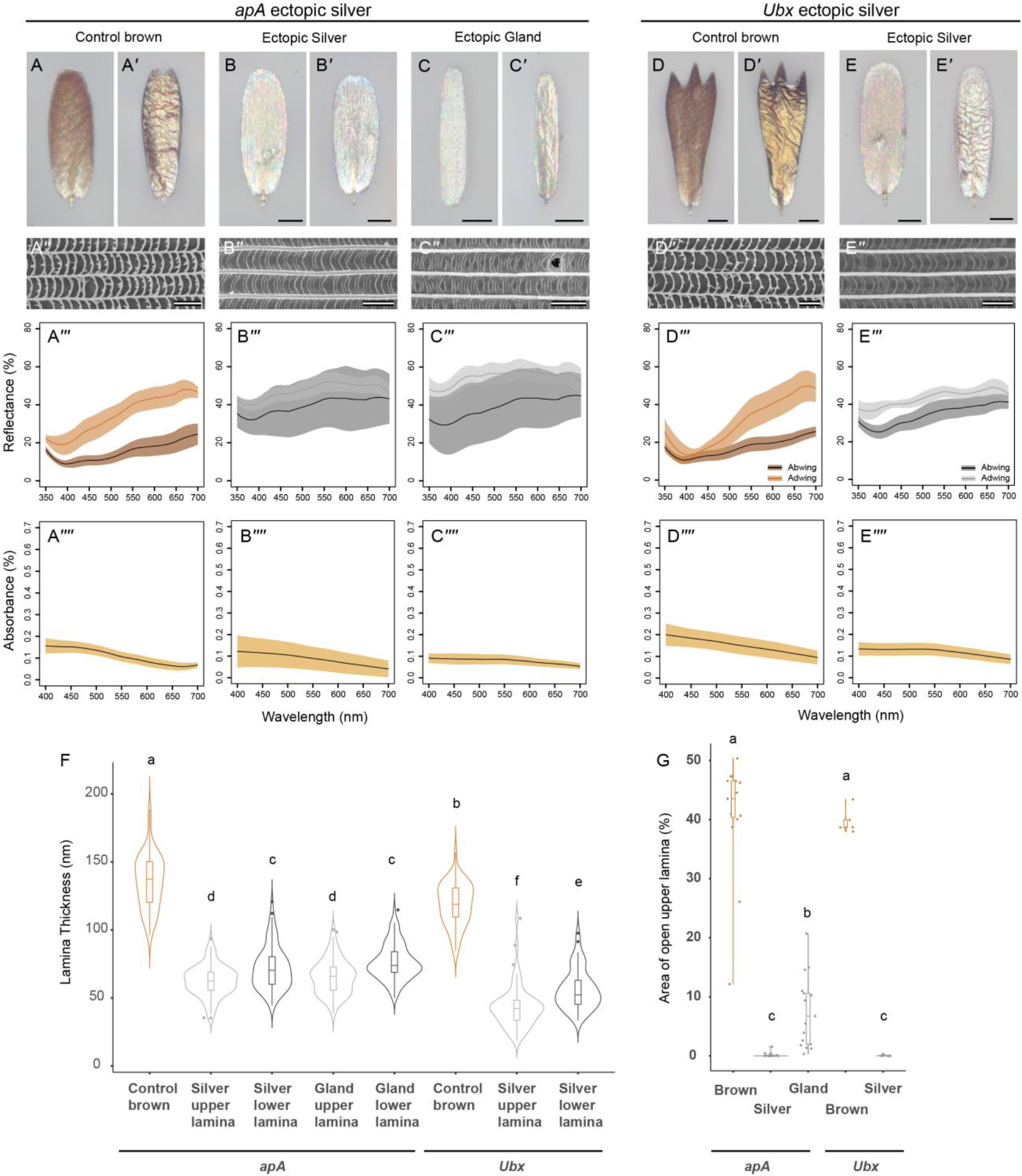
Characterization of ectopic silver scale ultrastructure and pigmentation in *apA* and *Ubx B. anynana* crispants. Optical microscopy images of the abwing (A-E) and adwing (A′-E′) surfaces, SEM images of the abwing surface (A′′-E′′), reflectance (A′′′-E′′′) and absorbance spectra (A′′′′-E′′′′) of the *apA* forewing control brown scales, ectopic silver and gland scales and the *Ubx* hindwing control brown scales and ectopic silver scales respectively. The control scales for these crispants are from the same wings as the ectopic scales, close to the mosaic crispant patches. (F) Violin plots of the upper and lower lamina thickness of the control brown scales and the ectopic silver and gland scales of *apA* and *Ubx* crispants. (G) Area of open upper lamina of the different control and ectopic scales. Boxplots show the median, inner and outer quartiles and whiskers up to 1.5 times the inter-quartile range. Means sharing a letter are not significantly different (Tukey-adjusted comparisons). Scale bars: A,A′-E,E′ are 20 µm and A′′-E′′ are 2 µm.

Structural modifications accompanying brown to silver scale transformations, in *apA* and *Ubx* crispants, were opposite to those seen when silver scales became brown. Ectopic silver scales gained a closed upper lamina and an enclosed air layer, reduced their lower lamina thickness, and lost pigmentation in comparison to control brown scales (Fig 4A′′-E′′, F, G). The upper lamina transformation was complete for the *apA* and *Ubx* silver scale subtypes, whereas many of the *apA* ectopic gland scales exhibited a partial transformation, having a significantly greater percentage of upper lamina open (Posthoc tests from LME, adjusted p-values <0.01; Supplementary Tables S1,2) (Fig 4G). The greater variability in the ultrastructural transformations and pigmentation of the *apA* ectopic silver scale subtypes explains the greater variability in the abwing reflectance spectra measured from these crispant scales (Fig 4B′′′, C′′′). All ectopic silver scale subtypes exhibited a significant decrease in lower lamina thickness when compared to the control brown scales (Fig 4F) (Posthoc tests from LME: adjusted P-values <0.0001; Supplementary Tables S3,4).

## Discussion and conclusion

### Metallic scales in *Bicyclus anynana* are undulatory broadband reflectors

We characterized five different metallic reflectors in *B. anynana* butterflies. Four out of the five scale subtypes are present only in males, associated with the androconial regions that produce and secrete sex pheromones, while the fifth, the coupling scales, are found in both sexes near the base of the wings. All five metallic scale subtypes investigated in this study produced broadband silver reflectance from an undulatory bilaminate structure achieved by the closure of the “open” windows present in typical wing scales. The air gap layer of varying thickness sandwiched between two thin chitinous laminas is spatially tuned to reflect different wavelengths of light, resulting in a specular broadband metallic color in the far-field due to additive color mixing. Such an ultrastructural modification involving the gain of a contiguous upper lamina leading to metallic coloration has been well-documented in multiple butterfly species across different families (21, 24), suggesting that the convergent evolution of metallic reflectors in butterflies can occur repeatedly with a few simple modifications to the basic scale architecture. Our modeling demonstrated that while butterflies can achieve broadband metallic colors by tuning just the lower lamina, the bilaminate three-layer system comprising a sandwiched air layer between chitinous laminas is substantially brighter, perhaps explaining its repeated evolutionary origin (24). Moreover, the varying thickness of the air layer was the most important structural parameter to produce a nearly flat (smoothed out) broadband reflectance, while the upper and lower lamina thicknesses were more optimized for enhancing the broadband reflectivity. Pigmentation levels were low overall but varied among the different silver scales along with slight variations in the total open area of the upper lamina. This intriguingly suggests that the amount of pigmentation may be directly impacting the formation of the upper lamina, as previously demonstrated in other scale subtypes of *B. anynana* (33). We were unable to characterize the exact pigment composition in these scales, but the absorbance spectra of the grey-silver scales suggests that the pigments are not purely melanins but could instead be a combination of melanins and additional pigments such as ommochromes (29).

### Each of five genes that led to gains and losses of metallic reflectance also changed pigmentation levels and scale ultrastructure dimensions

We discovered that disruptions to the five genes investigated (*Antp, dsx, apA, Ubx and optix*) lead to simultaneous change in three scale features: presence or absence of an upper lamina, pigmentation levels, and lower lamina thickness. Transformation of metallic silver scales into brown, non-reflecting scales in *Antp, dsxM, apA* and *optix* crispants involved loss of an upper lamina, gain in pigmentation levels, and an increase in lower lamina thickness. This was generally true except in the case of the grey-silver scales of the hindwing where transformation to brown scales led to a decrease in pigmentation. Conversely, gain of silver reflectance in scales of *apA* and *Ubx* crispants was accompanied by the appearance of a continuous upper lamina, a decrease in pigmentation levels, and a decrease in the thickness of the lower lamina. The breadth of variation seen in individual crispant brown scales and ectopic silver scales was potentially due to incomplete knockouts, and variation in the protein function of each mutant allele. This was especially the case of *apA* crispants, possibly due to the incomplete knockout of these genes in individual scales cells. The gain and loss of an upper lamina with concurrent changes in the thickness of the lower lamina suggests a possible conservation of total chitin production within a cell, but variable deposition between the two laminas. There is some evidence for this in the forewing silver and gland scales but not in the hindwing silver scale subtypes. The summed thicknesses of the two laminas in the wildtype forewing silver scales and ectopic silver scales was similar to the lower lamina thickness of the forewing control brown scales and crispant brown scales. Though there is some indication of conservation of chitin production within a single cell, testing this proposition in the future will require finer measurements of chitin production within different scale color types. Further, Matsuoka and Monteiro (33) speculated that downstream effector genes such as *yellow* could affect cuticle polymerization around crossribs to create windows because *yellow B. anynana* crispant black and brown scales exhibited an upper lamina covering the windows. Given the low amounts of pigmentation seen in the different silver scale subtypes, it is possible that the gene *yellow* is not highly expressed in silver scales, mirroring *yellow* crispant scales. *yellow* would then be a potential downstream target, that would be repressed by the silver scale differentiation program. This hypothesis could be tested by measuring levels of *yellow* expression in the different colored vs silver scales of *B. anynana*, as well as in crispant individuals during development.

### *Antp* and *optix* are high-level regulators in the differentiation of silver scales, affecting both scale color and shape

Investigation of the mutant brown scales in *Antp* crispants uncovered an important role for this gene in determining a rounded scale morphology. Scales on the posterior ventral forewing and anterior dorsal hindwing of both sexes of *B. anynana*, a region where Antp is strongly expressed (Supp Fig S7), normally have rounded distal ends, appearing silver in males and shades of brown in females. In *Antp* crispants, the rounded scales developed serrated distal tips in both sexes, concurrent with the transformation of silver to brown color in males only. These crispant brown scales with serrated edges resembled scales usually found on the rest of the ventral forewing (Supp Fig S6A, B) or dorsal hindwing (Supp Fig S6C, D), indicating that *Antp* is necessary for converting a serrated distal scale edge into a rounded scale. In contrast, while both *Antp* and *dsxM* affected scale color, knockout of *dsxM* did not affect scale shape. Thus, though both genes are expressed in a trait-specific manner, *Antp* affects aspects of scale color and shape while *dsxM* has more restrictive transformations of color only. The exact hierarchy of interaction between *Antp* and *dsxM* in silver scale differentiation remains to be tested. Both genes could be acting in parallel or *Antp* could be upstream of *dsxM* (Fig 5). Similar to *Antp, optix* also exhibits high levels of expression in a trait-specific manner i.e., in the coupling scales, and is necessary in determining both color and shape of the coupling scales. These data suggest that *Antp* and *optix*, are important regulators in determining the differentiation of various silver scale subtypes. Both genes are highly expressed in the silver scale cells and affect multiple aspects of their development including the color and the morphology. Both genes are necessary for silver scale development but are not sufficient given that both *Antp* and *optix* have other expression domains on the wing like the eyespot centers (*Antp*) or the gold ring (*optix*), which aren’t silver. Furthermore, the conserved expression and function of *optix* in coupling scales in several other butterfly species implicates a conserved role for this gene in the development of trowel-shaped coupling scales (37, 38).

**Figure 5:**
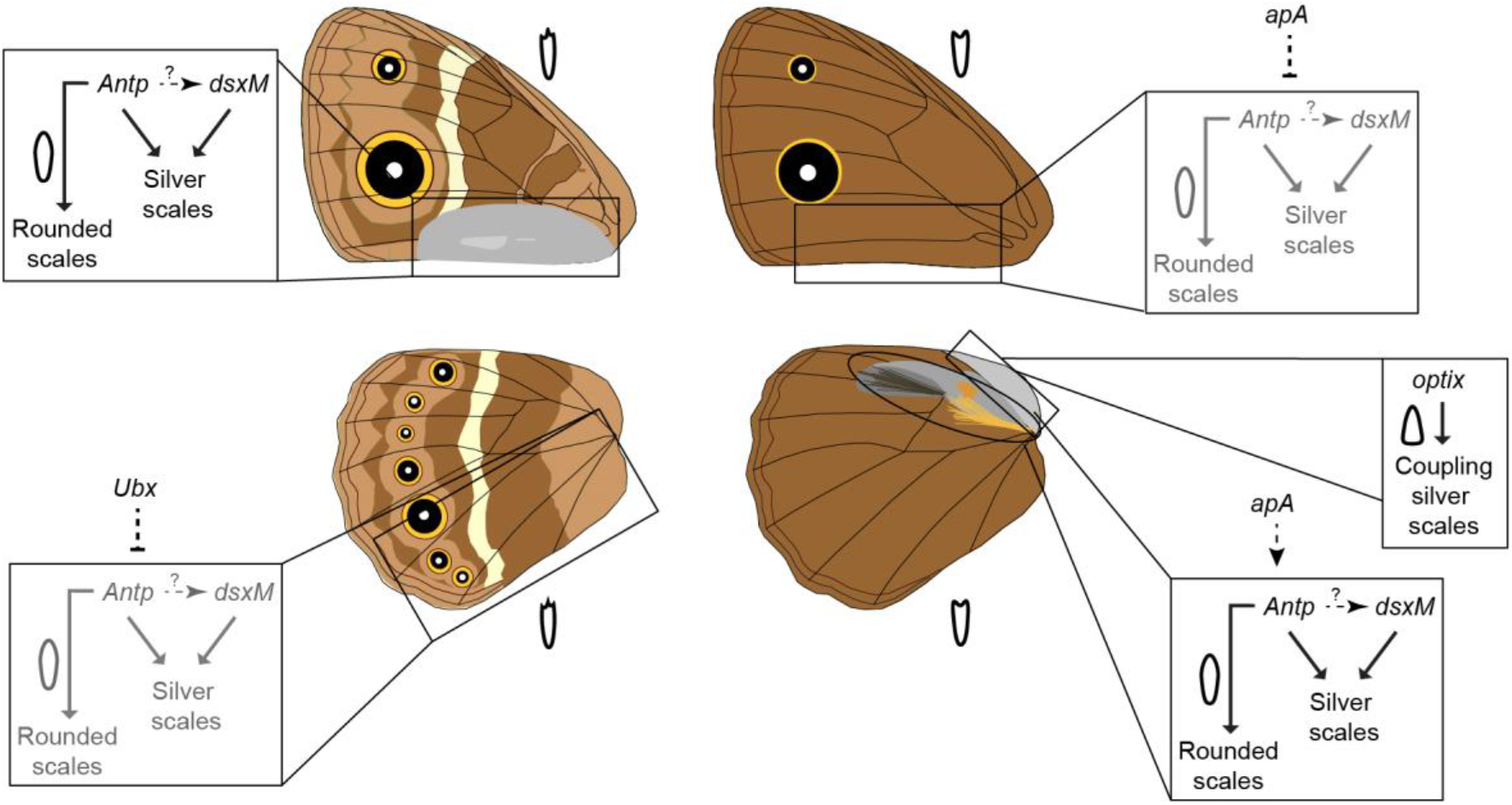
Functions and interactions of genes involved in the differentiation of different silver scale subtypes across wing regions of a male *B. anynana*. For each wing region, the genes involved in differentiating silver scales, their interactions and functions are indicated within boxes. The homeotic selector genes *apA* and *Ubx* are outside the boxes. Their relation to the differentiation of silver scales is indicated by dotted lines because a direct or indirect interaction between *apA/Ubx* and the genes within the boxes remains unknown. The scale schematics inside the boxes represent the scale shape when the ‘rounded scales’ part of the network is active (dark lines). Repressed parts of the network are in grey. The scale schematics outside the boxes, next to each wing surface, reflect the scale shape generally found on that surface otherwise.

Interestingly, the metallic silver scales on male forewings and hindwings fall into two populations with different aspect ratios: the shorter silver scales covering most of the region and the longer, thin gland scales that cover only the underlying pheromone glands (41). The spatially defined occurrence of the long, thin gland scales is likely specified by the spatially restricted expression of some unknown gland-specific gene, downstream of DsxM in the silver scale differentiation program. In addition, the difference in levels of Antp protein expression between the silver and grey-silver scale regions suggests that varying thresholds of this protein are functionally relevant in the two scale subtypes.

The unusual ridge and crossrib orientation, almost orthogonal to the normal orientation, seen in a few *Antp* crispant brown scales, hinted at a role for this gene in ridge patterning during the early stages of scale development. In developing scales, the positioning and orientation of ridges is determined by the spacing of F-actin filaments that emanate from the base of the scale, parallel to the scale’s long axis (42, 43). The disrupted and abrupt change in orientation of the ridges and their corresponding crossribs in the *Antp* crispant scales suggests a potential disruption of the orientation of F-actin filaments. Based on the distal domain of occurrence of the unusual ridge patterns, along with the rounded to serrated distal edge change seen in *Antp* crispants, we speculate that *Antp* regulates the distal expression of an unknown factor, possibly a cytoskeletal component, that determines scale morphology and ridge/crossrib orientation. Hox genes such as *Abd-B* have been identified to activate cytoskeletal components like cadherins in the development of posterior spiracles of *Drosophila* (44). Furthermore, *Ubx* directly regulates cytoskeletal components such as actin and contributes to cytoskeletal reorganization in haltere cells (45). In that context, our results identify a novel post-embryonic role of a Hox gene in the determination of the shape of a single cell. How *Antp* directs scale shape and ridge orientation, and which genes are its downstream targets, remains an exciting avenue for future work.

### Homeotic genes and the differentiation of silver scales

As predicted in our hypothesis, transformed scales investigated in the *apA* and *Ubx* crispants exhibited changes in both color and shape that resembled the wildtype silver or brown scales in that region. Given that both genes have broad expression domains in the dorsal wing compartment (*apA*) (35) or hindwings (*Ubx*) (36) of *B. anynana*, these genes may not constitute part of the silver scale differentiation program. However, these selector genes are likely regulating the wing and surface-specific expression of genes like *Antp, dsx* and *optix* by binding to trait-specific enhancers of these genes in a combinatorial manner with other transcription factors. Searching for *apA* or *Ubx* binding sites in regulatory regions around *Antp* or *optix* and functionally testing them via targeted deletions of these regions is one way of answering this question in the future. In any case, *Ubx*, in particular, appears to repress the differentiation of silver scale on the hindwings of *B. anynana* while *apA* has either a repressive or activating function for silver scale differentiation on the forewings and hindwings, respectively (Fig 5). The direct or indirect nature of these interactions remains unknown.

In conclusion, this study identified the structural and photonic origin of broadband silver coloration by comparing silver scales in WT and crispants for five genes in the butterfly *B. anynana*. We identified *Antp* and *optix* as important high-level regulators in the differentiation of silver scales and identified a novel role for these genes in determining the shape of single cells.

## Materials and methods

### Crispant individuals and sampled scale subtypes

*B. anynana* crispants for *apterous A* (*apA*) (35), *Ultrabithorax* (*Ubx*) (36), *doublesex* (*dsx*) (34) and *Antennapedia* (*Antp*) (36) were previously generated in our lab by CRISPR-Cas9 mediated gene editing. Crispants for *optix* were generated by injections of two guide RNAs targeting the *optix* coding sequence (46). Five different silver scale subtypes were sampled, namely, the silver and the gland scales on the androconial patch of male ventral forewings, the silver and grey-silver scales near the androconial patches of male dorsal hindwings and the coupling scales near the base of the dorsal hindwing in females (Fig 1A).

### Scanning electron microscopy

Five to six scales from different regions of the crispant individuals (Fig 1A) were individually mounted onto carbon tape and sputter coated with gold using a JFC-1100 Fine Coat Ion Sputter (JEOL Ltd. Japan). Images were obtained using a JEOL JSM-6510LV scanning electron microscope (JEOL Ltd. Japan). For the wildtype, *apA* and *dsx* crispants, three different individuals were sampled for each case. Two individuals were sampled for *Antp* and *optix* crispants while only one crispant was obtained for *Ubx*.

### Percentage area of open upper lamina measurements

From the SEM images, the percentage of area of open upper lamina was calculated using ImageJ 1.52a (Java 1.8.0_112) (47). A 20 µm^2^ (10 µm^2^ for gland scales) region of interest was defined approximately at the center of the scale and the region outside cleared. A thresholding was applied based on the values of the bright (ridge and crossribs) and dark (windows) regions of the SEM. The dark areas were selected and added to the ROI Manager in ImageJ using the Analyze Particles function. The selected regions were combined using the OR function in the ROI Manager. The combined area of the open upper lamina within the defined regions was measured and converted into a percentage.

### Focused ion beam scanning electron microscopy (FIB-SEM)

FIB-SEM was used to measure the lower lamina, upper lamina, and air gap thicknesses for all scale subtypes in our analysis and corrected for tilted perspective (measured thickness / sin 52°). Briefly, samples were prepared by sputter-coating with platinum to increase conductivity. The scales were milled using a gallium ion beam on a FEI Versa 3D with the following settings: beam voltage −8kV, beam current −12pA at a 52° tilt. Image acquisition was performed in the same equipment with the following settings: beam voltage −5kV, beam current −13pA. Milling was done at the center of each scale. Thickness measurements were done in ImageJ. For each scale subtype, ten measurements were taken per scale with 3-16 scales sampled from 1-2 individuals. Measurements were made along most of the lower lamina which is uniform, excluding the region around the ridge base, where the thickness is highly variable.

### Optical imaging and UV-VIS-NIR microspectrophotometry

Light microscope images of individual scales were recorded using the 20X lens of a uSight-2000-Ni microspectrophotometer (Technospex Pte. Ltd., Singapore) and a Touptek U3CMOS-05 camera. Scales were individually mounted on a glass slide or in a refractive index matching medium (clove oil) and multiple images at different focal planes (z-stack) were obtained. Stacking was done in Adobe Photoshop v 22.5.1 (Adobe, California, USA).

Normal-incidence UV-VIS-NIR reflectance spectra of scales were acquired using the same microspectrophotometer setup but with a 100x objective. Spectra with usable range between 335-950 nm were collected using a high NA 100x objective from a ∼2 μm sized spot (100 ms integration time, 10x averaging) and calibrated using an Avantes WS-2 reference tile made of white diffuse polytetrafluoroethylene. Individual scales were mounted on a black carbon tape and illuminated with a Mercury-Xenon lamp (ThorLabs Inc., New Jersey, USA). Measurements were taken from both abwing and adwing surfaces. For each scale subtype, measurements from five to ten individual scales from one individual were averaged. Absorbance spectrum was measured for individual scales immersed in a refractive index matching liquid (clove oil) using a 20X objective. Six to eight individual measures from one individual were averaged for each crispant type and wildtype. Analysis and spectral plots were done in R Studio 1.4.1106 with R 4.0.4 (48) using the R-package *pavo* (v 2.7) (49).

## Acknowledgements

We thank Yuji Matsuoka for providing us the *Antp* and *Ubx* crispants and Emilie Dion for helpful discussions on the statistics. We are grateful to Dr. Robert Reed for providing us the Optix antibody. We thank Sree Vaishnavi Sundararajan and Gianluca Grenci (MBI) for access and help with SEM, and the Pennycook group (MSE) for use of FIB-SEM. This research was supported by the National Research Foundation (NRF) Singapore under the Competitive Research Programme (NRF-CRP20-2017-0001 Award) and the National University of Singapore.

## Author Contributions

A.P and A.M conceived and designed the study. A.P, C.F and V.S collected the spectral measurements. A.P and C.F collected the SEM data and C.F collected the FIB-SEM data. V.S performed the theoretical modeling. A.P analyzed all the data and did the Antp immunostainings. T.D.B performed the Optix immunostainings and the *optix* knockout experiments. A.P wrote the manuscript with inputs from all the authors.

## Notes

https://github.com/evolphotonics/bbandAgmodel

### Competing Interest Statement

The authors have declared no competing interest.

### Summary of Updates

Author list and affiliations updated; Involvement of optix in silver scale development in Bicyclus anynana updated; Advantages of a bilaminate scale structure for silver color production highlighted; silver scale differentiation program and roles of different genes updated

