## Supplement for "*Antennapedia* and *optix* regulate metallic silver wing scale development and cell shape in *Bicyclus anynana* butterflies"

This file includes:

- Supplementary Methods
- References for SI
- Supplementary Figures S1-S8
- Supplementary Tables S1-S4
- Supplementary Table S5 – source data in an excel sheet

**Supplementary Methods**

Theoretical modeling

To model the theoretical reflectance of scale ultrastructural elements, both individually as well as in various combinations, we used the freely available FreeSnell software (A. Jaffer, http://people.csail.mit.edu/jaffer/FreeSnell), a thin-film optical simulator implemented in the SCM Scheme language. As inputs to the model, we used the upper and lower lamina and air gap thickness measurements (see Fig 1I, and Supplementary Table S5 – source data) from FIB-SEM cross-sectional images of the various scale types. The calculations were first performed in a hierarchical fashion – varying only upper lamina, air gap or lower lamina thickness, then both upper and lower lamina thicknesses, and lastly all three simultaneously. Our modeling suggested that only 3-layer models that include a varying air gap layer were able to explain both the broadband nature and the overall brightness of the measured scale reflectivities, as compared to 3-layer models with constant air gap thickness but varying upper and/or lower lamina thicknesses or 1-layer models with just an upper or lower lamina (see Supp Figs S3, and S4).

We used the following variational method to generate the mean theoretical reflectance (with error bars) for the five different silver scale types reported in Figs. 1B‴-F‴, and Supp Fig S2. For each scale type, we averaged 200 spectra generated using a 3-layer model – where only the air layer thickness parameter is varied using a set of $N = 200$ values specified using a normal distribution with the measured mean and standard deviation (see Fig 1I, and Supplementary Table S5 – source data), while the upper and lower lamina thicknesses were constant and set to their respective measured mean values. We plot the theoretical abwing spectra in Figs. 1 and S2 after accounting for the short-wavelength absorption due to pigments (1B⁗-F⁗). Final predicted abwing reflectance were obtained by multiplying the theoretical reflectance by the corresponding percentage pigmentary transmittance ($T$, given by $10^{-A}$, where $A$ is the measured absorbance of the scale type).

In order to perform the systematic parametric sweeps presented as heatmaps in Supp Fig S4, for each scale type, two of the three structural parameters were held constant and set to their measured mean values (see Fig 1I, and Supplementary Table S5 – source data), while the third layer thickness was allowed to vary over a range of values in small discrete steps (0 to 2 μm for air gap in 20 nm steps, 0 to 500 nm in 5 nm steps for both lower and upper lamina). Further at each step, we used the variational method described above to generate and average over the spectra corresponding to $N$ = 100 values specified by a normal distribution centered on the step value (*i.e.* mean), and a standard deviation scaled by the coefficient of variation of the corresponding layer's measurement (step standard deviation = step mean * measured standard deviation / measured mean; for instance, the standard deviation corresponding to a mean forewing silver upper lamina thickness of 500 nm works out to be 500 nm * 8 nm / 64 nm = 62.5 nm). Here, we used $N$ = 100 in contrast to the $N$ = 200 used earlier, in order to save computational time, and having verified that they both produced near identical mean spectra. At each step, we also averaged the mean spectra over the entire wavelength range and plotted this single value (mean broadband reflectivity, $R$) as a function of the varied thickness parameter as an inset (Supp Fig S4).

Random value generation and averaging of spectra were performed using custom code written in *bash* and *awk* scripting languages. We used the stackplot functionality of *pavo* package (v 2.4) (1) in R (v 3.6.3) (2) to produce the extrapolated heatmaps in Supp Fig S4. In order to average over any spatial differences in pigment deposition, the absorbance data were measured using a 20X objective that does not extend into the UV. The absorbance measurements were therefore extrapolated into the UV using local polynomial regression with the help of *loess* (span of 0.2) and *predict* functions in R (2). Lastly, the layer thickness measurements plotted in Supp Fig S3C were made on a FIB-SEM cross-section image using the BAR plugin 1.5.2 (doi: 10.5281/zenodo.495245) in FIJI (v 1.5.3) running on a Linux platform. Briefly, a ~1.7 μm wide area region of interest (ROI) was selected, air region in the exposed plane masked and thresholded to convert into a binary image, and then filtered (3x3 median) to remove noise. A vertical line ROI was drawn at the left end of the image ROI and translated by 6 pixels (or ~20 nm) along the width of the image to the other end. At each of these 82 loci, the corresponding line profile was plotted. A horizontal line ROI was plotted at the half-height of this plot profile and subsequently its line profile plotted, in turn. The position of the air-chitin interfaces were found using the "Find Peaks" functionality of the BAR plugin as applied to this second-order line profile, using default parameters. These tasks were automated using a custom macro. The difference between consecutive minima gave the relevant layer thicknesses (see Supplementary Table S5 - source data). The color swatch at each of the 82 line profile locations were generated by FreeSnell as 64x64 pixel color squares by invoking the color-swatch function that uses the CIE illuminant D65.

The entire set of code for performing the broadband reflectance simulations is publicly available in the following GitHub repository: https://github.com/evolphotonics/bbandAgmodel.

Trichromat insect visual modeling

In order to understand how butterflies could perceive the theoretical spectra produced by the various models (Fig S3B), we performed visual modeling using *pavo* (v 2.4) (1) in R 3.6.3 (2). The relative stimulation (quantal catches) of the three photoreceptors (*s*, *m*, and *l*) and the corresponding colorimetric parameters in a trichromat color space (with chroma or saturation, the magnitude of the position vector given by the distance from the achromatic centroid of the color triangle) were computed and plotted with the Vinegar Fly (Diptera: *Drosophila melanogaster*) spectral sensitivities and default settings (homogeneous illuminance). This was the closest available visual model specified in *pavo* to *Bicyclus anynana* butterflies, in terms of phylogenetic relatedness (Diptera is sister to Lepidoptera) and the overall similarity of their visual sensitivities (3).

Statistical analysis

Statistical analyses were performed in R Studio 1.4.1106 with R 4.0.4 (2). The differences in mean thickness or mean percent area of open upper lamina among the different crispant types and wildtype were analyzed using a linear mixed-effects model (LME) that allows for fixed and random effects. Due to the hierarchical nature of the datasets, where multiple measurements were taken from each scale and multiple scales were analyzed for each individual, we used a LME with the crispant type as the fixed factor, and scale nested within individual as a random factor. LME was run using the nlme package (v 3.1.152) (4). In addition, due to the violation of the homogeneity of variance in the measurements among different crispant types or wildtype, we allowed for different variance structures for each type using the ‘varIdent’ function in the nlme package. Different models were compared using the anova function and the best model was selected using AIC as a criterion. Adjusted P-values for different pairwise comparisons were obtained by a posthoc analysis using the multcomp (v 1.4.17) package (5) and Tukey contrasts. Outcomes of the LME tests and the adjusted P-values for multiple comparisons for all the tests are given in Supplementary Tables 1-4.

Antp and Optix Immunostaining

The Antp primary antibody used was the same as (6). anti-Antp 4C3 was deposited to the Developmental Studies Hybridoma Bank by D. Brower. The secondary antibody was an Alexa Fluor 488-conjugated goat anti-mouse antibody (Jackson ImmunoResearch Laboratories, Inc.). The Optix primary antibody (from rat) was a gift from Robert D. Reed and was the same as (7). The secondary antibody was an anti-rat AF 488 (Invitrogen, #A-11006).

Pupal wing tissues were dissected in phosphate-buffered saline (PBS) and transferred to fix buffer at room temperature (0.1M PIPES pH 6.9, 1mM EGTA pH 6.9, 1% Triton X-100, 2mM $MgSO_4$). Wings were fixed in 4% formaldehyde (added to the wells) at room temperature for 30 minutes, then washed five times with PBS. Blocking was done overnight at 4 ˚C in block buffer (50mM Tris pH 6.8, 150mM NaCl, 0.5% IGEPAL, 1mg/ml BSA). The wings were then incubated in primary antibody (1:100 of 1:1 anti-Antp:glycerol solution or 1:3000 of anti-Optix) in wash buffer (50mM Tris pH 6.8, 150mM NaCl, 0.5% IGEPAL, 1mg/ml BSA) at room temperature for 1 hour, washed with wash buffer four times and then incubated with secondary antibody (1:500) for 30 minutes at room temperature. Samples were washed to remove the secondary antibody and incubated with DAPI for 5-10 minutes, followed by further washing. Wings were imaged on an Olympus FLUOVIEW FV3000 confocal microscope.

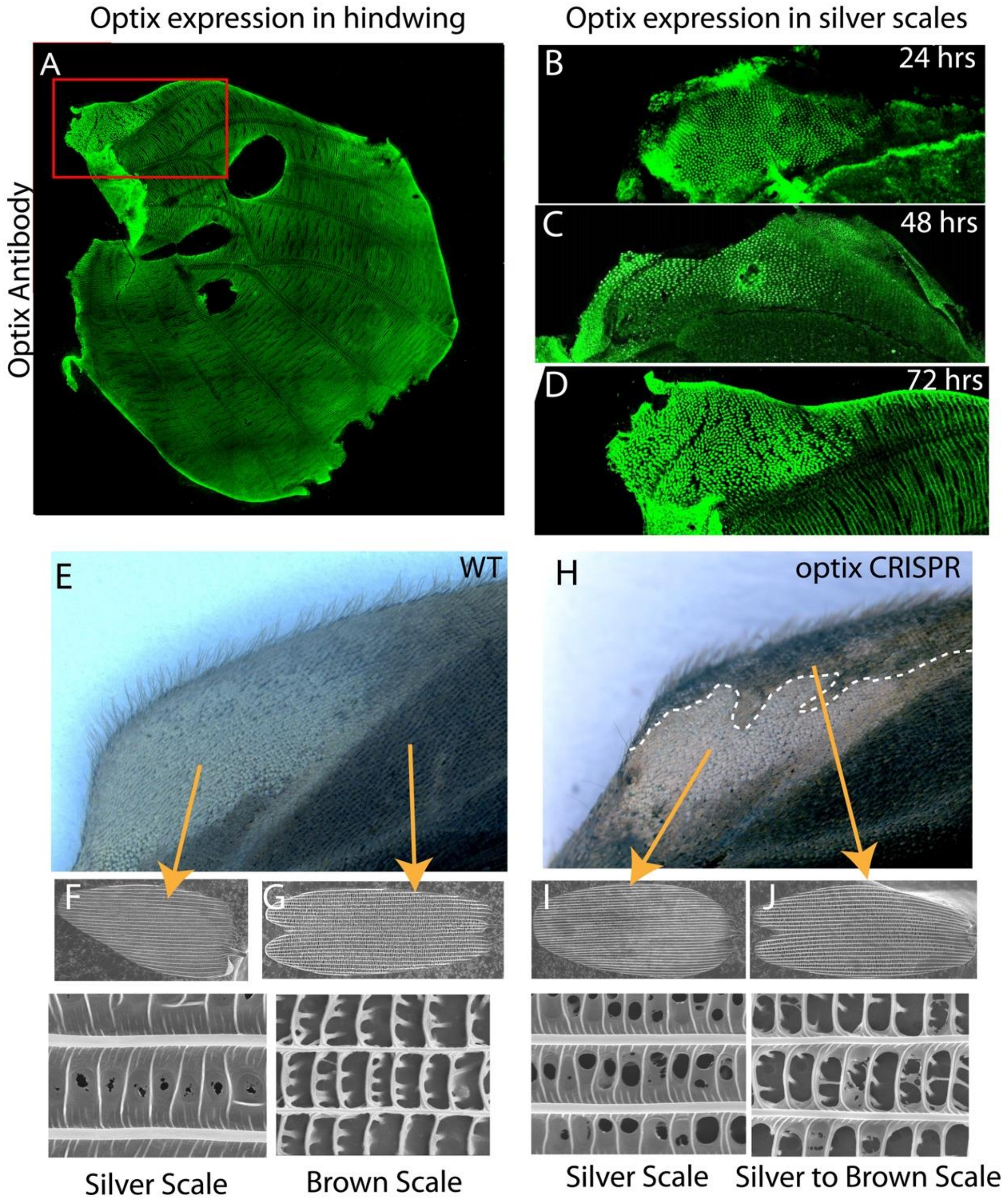


**Supplementary Figure S1: Optix protein expression in the pupal hindwings of *Bicyclus anynana* and crispant phenotypes in the silver scale region**. (A) Hindwing showing expression of Optix proteins in the future silver scales. Expression of Optix at (B) 24 hrs, (C) 48 hrs and (D) 72 hrs pupal hindwing. (E) WT hindwing (F, G) Ultrastructure of a WT silver coupling scale and a WT brown scale. (H) *optix* crispant hindwing. *optix* CRISPR results in the conversion of silver coupling scales into brown scales. (I, J) Ultrastructure of a silver coupling scale and a silver coupling scale converted into a brown scale from an *optix* crispant hindwing. The crispant brown scale structure resembles that of a WT brown scale with open windows.

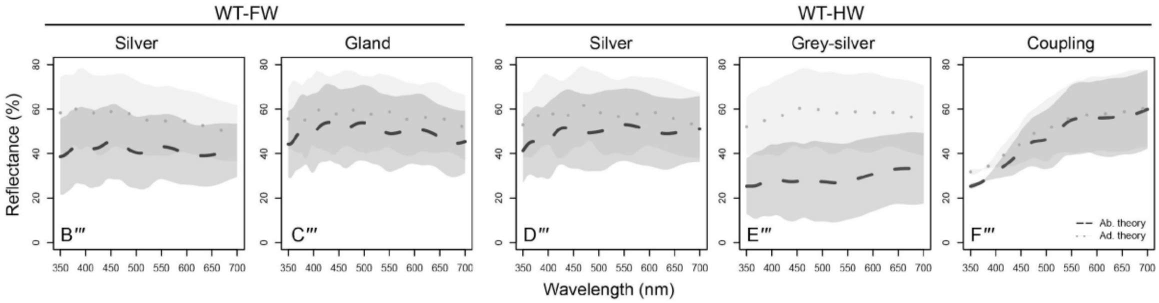


**Supplementary Figure S2: Theoretically modeled reflectance spectra for the five different wild-type silver scales of *Bicyclus anynana***. The abwing (dashed line) and adwing (dotted line) spectra are reproduced here from Fig 1B‴-F‴, along with shaded areas representing one standard deviation from the mean (dashed or dotted lines). The abwing spectra include the corresponding pigmentary absorption shown in Fig 1B⁗-F⁗. The theoretical modeled spectra which were computed with respect to a specular standard have been offset vertically (+30%) so that they are comparable to experimental reflectance measured relative to a diffuse standard (Fig 1B‴-F‴). See supplementary methods for details.

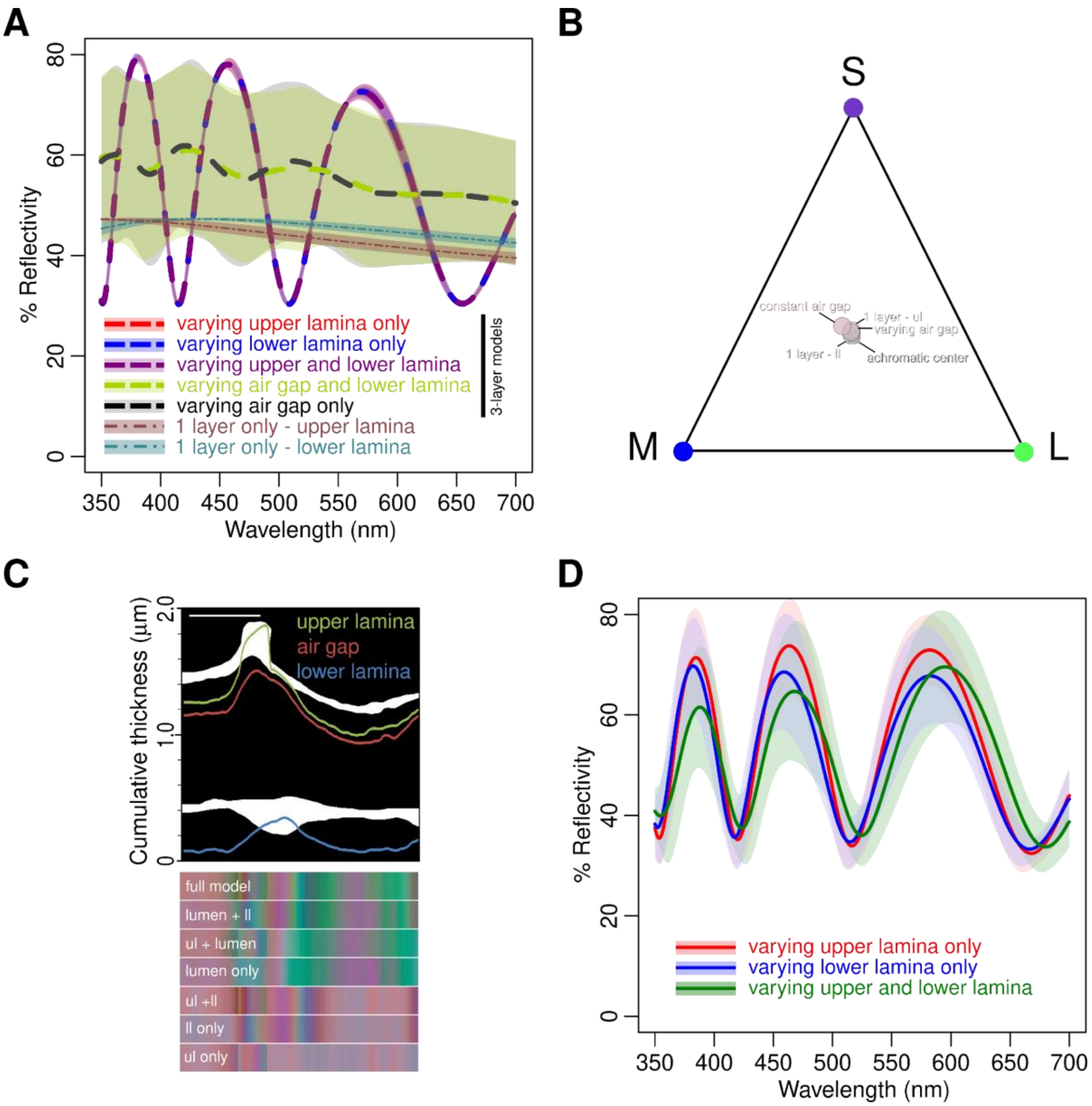


**Supplementary Figure S3: Hierarchical optical modeling of the broadband metallic color production in forewing silver scales of *B. anynana*. A**) Mean modeled spectra are plotted for the full 3-layer models (dashed curves) and 1-layer models (dash-dot curves) using the variational method (see supplementary methods for details) with shaded areas representing 1 standard deviation. For 3-layer models, either one or two specified (see legend) parameters are varied, while keeping the other parameter(s) constant. An enclosed air gap layer (lumen) of variable thickness is necessary to produce a bright, relatively flat broadband (achromatic) reflectance, compared to the rather chromatic sinusoidal spectra when air gap thickness does not change or the more broadband (flat) but much lower (duller) reflectance from a 1-layer model with only a varying upper or lower lamina. Note, the 3-layer models with a varying upper (red curve) or lower lamina (blue curve) essentially overlap, occluding the former from view. **B**) The mean modeled reflectance in **A** plotted according to a *Drosophila* trichromat visual model further affirms that only nanostructural models with varying air gap layer (grey circles) produce broadband colors that are closer to the achromatic centroid (light grey square), compared to models where the air gap thickness is constant (pale pink circles). **C**) In contrast to the spectra in **A**, which were modelled using the mean and variance of lamina and air gap thickness measurements (Supplementary Table S5) made in the closed window areas (away from the ridges), here we model the color and spectra directly from a silver scale FIB-SEM cross-section, thereby accounting for the thickening of the laminas near the ridges. Top panel: A binarized (chitin – white, air – black) region of interest (ROI) from a scale FIB-SEM cross-section overlaid with the corresponding cumulative thickness profiles of the three layers (colored lines). Scale bar for *x*-axis (500 nm) differs from *y*-axis as the latter has been corrected for perspective foreshortening in FIB-SEM measurements. Bottom panel: 82 conjoined color swatches for each of the 7 possible hierarchical models where 1, 2 or all 3 nanostructural layers are varied are shown at ~20 nm intervals (i.e., every 6 pixels) along the width of the ROI. Models incorporating the varying air gap layer are closest in hue to the full 3-layer model, whereas the effects of the laminas are most pronounced at or near ridges, far from inter-ridge regions where they have nearly uniform thicknesses. The blue-green and purple-pink hues in between ridges here correspond well with similar color highlights seen in high magnification images of the scale in between ridges (Fig 1B-F). **D**) Averaging the spectra corresponding to the 82 color swatches in **C** for the 1- and 2-layer lamina models understandably shows more variance as compared to **A** ($N$ = 200), however, the resulting spectra here are still similarly chromatic and sinusoidal. When laminar thicknesses vary more at the ridges, interference from upper and lower lamina reduces the intensity at shorter wavelengths. This perhaps explains a similar reduction in the measured reflectivities at short wavelengths, relative to our simplified theoretical predictions that does not account for the thickening at and around ridges (Fig 1B‴-F‴, and Supp Fig S2). Abbreviations: ul – upper lamina, ll – lower lamina.

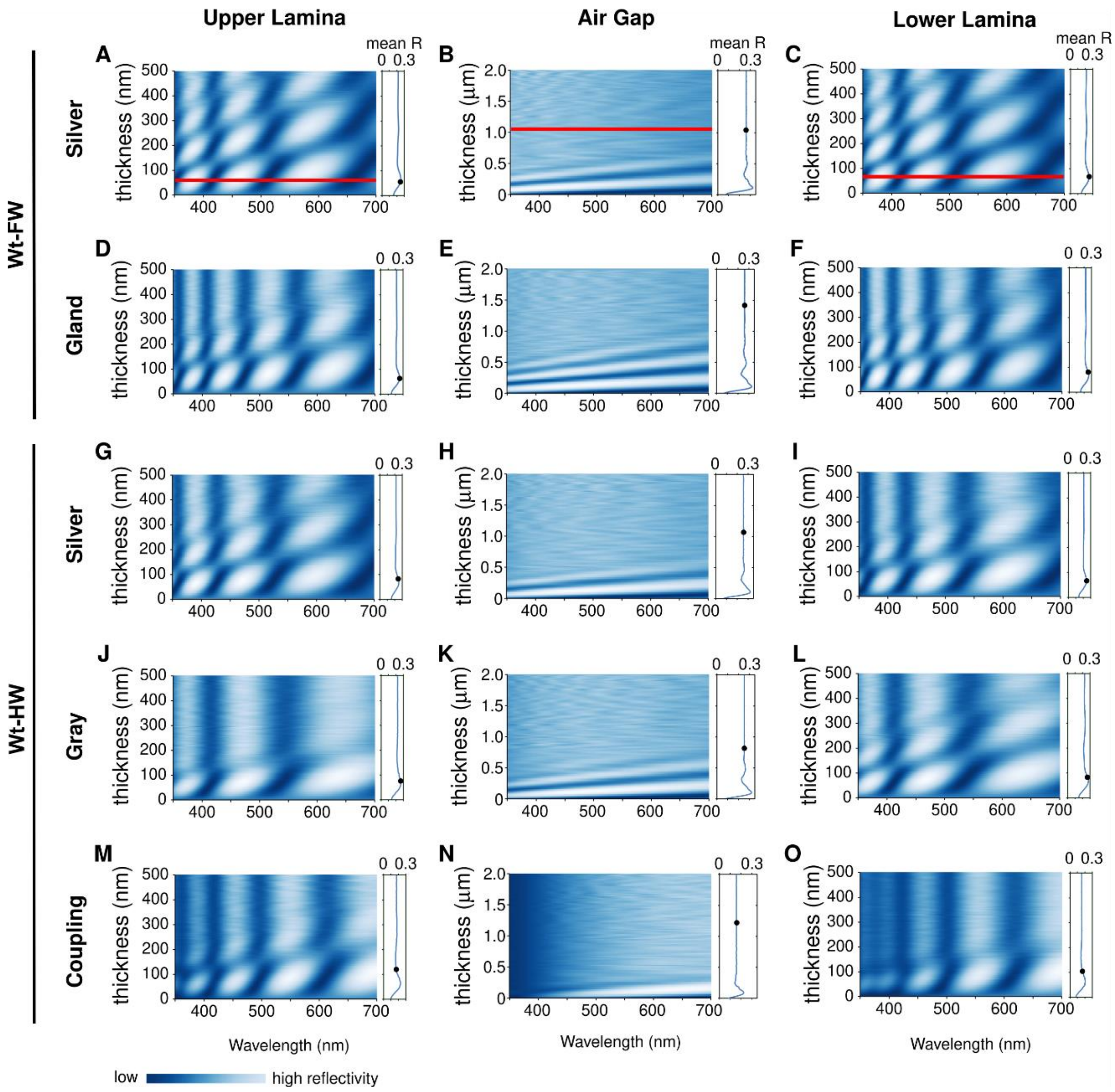


**Supplementary Figure S4: Simulations over the parametric space of the five different wildtype silver scales of *B. anynana* to understand how changing one layer thickness at a time affects broadband color production.** Theoretical reflectivities (relative to a specular standard) modeled by systematically changing the layer thickness of upper lamina (first column), air gap (second column) or lower lamina (third column), while keeping the other two parameters constant, are plotted as heatmaps. The heatmaps are top-views of a complex 3-D dataset of 100 spectra extrapolated across the range of thickness values (with wavelength along *x*, thickness along *y* and reflectivity/intensity along *z*), so that the peaks (high intensity) in the spectra are shown in shades of white and dips (low intensity) in navy blue. For reference, the reflection spectra corresponding to the thick red lines in **A**-**C** are as plotted in Fig. S3A (red, blue and black dashed curves). The inset plots on the right show the mean broadband reflectivity, *R*, averaged over the entire wavelength range at each thickness value and the black dot denotes the corresponding measured layer thickness for that specific scale type (see Fig 1I, and Supplementary Table S5 – source data). *R* is largely constant across the thickness range but shows a small peak at the lower end of the range. The observed lower and upper lamina thickness for all silver scales except coupling seem to lie at or very close to this peak, *i.e.*, they seem to be optimized for maximal broadband brightness. The air gap thicknesses are, however, not optimized for maximal brightness (at the lower end of the range), but rather thick enough to ensure the reflectivity is achromatic and broadband (i.e., uniform or flat). Varying only upper (left column) or lower lamina (right columan) thicknesses even upto 500 nm shows that the spectra still remain sinusoidal and chromatic because of the pronounced peaks (strong white areas). Only by varying air gap thickness (middle column), do we obtain nearly uniform broadband reflectivity above a certain thickness threshold (diffused patterns in column B). The coupling scales, which are the least pigmented among any of the silver scale types, are a natural experiment that illustrates the effect of increasing the upper and lower lamina thickness beyond their optimal values. Similar to the effect of the thickened lamina near the ridges (Supp Figs. S3C, and D), the coupling lamina thicknesses, which are above the optimum, reduce the intensity at shorter wavelengths (notice the darker blue color in Figs. S4M-O at these wavelengths), analogous to the effects of short-wavelength absorbing pigments (see Fig 1B''''-F'''').

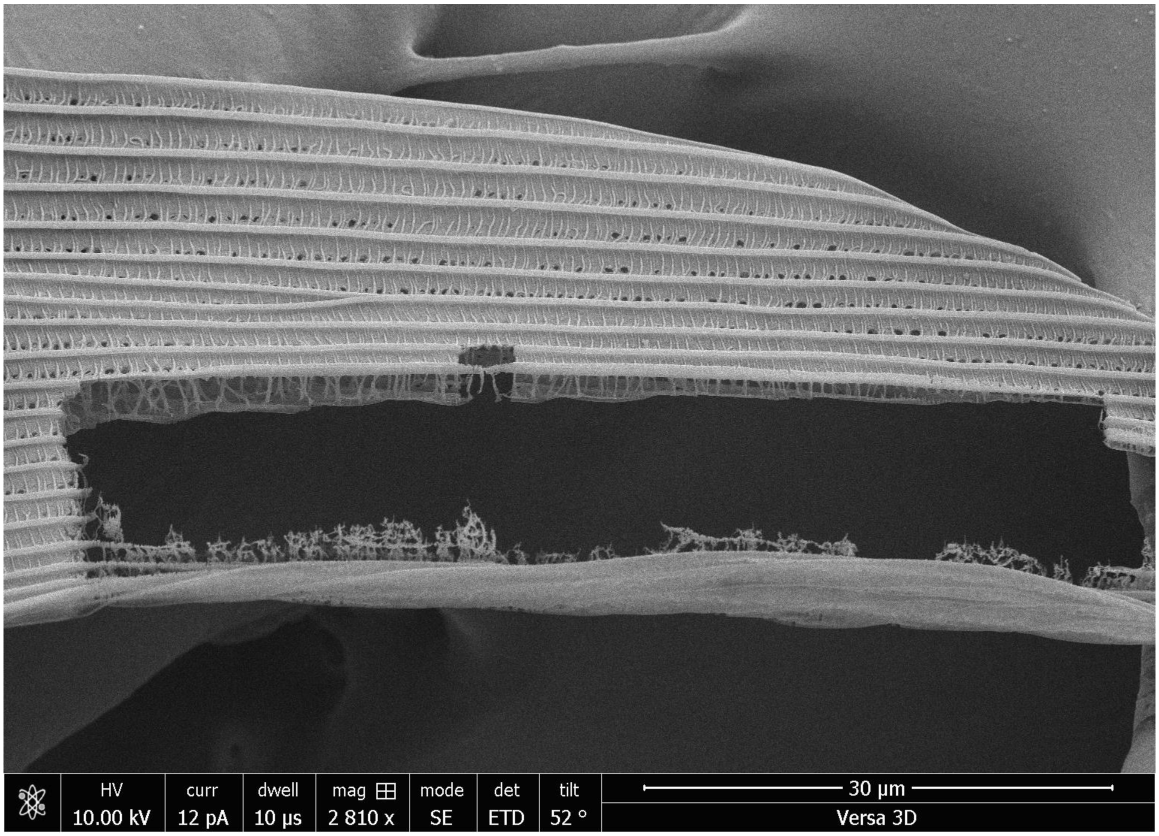


**Supplementary Figure S5: Longitudinal cross-section of a wildtype coupling scale of *Bicyclus anynana*.** Proximal base of the scale is to the left. The thickness of the lower lamina decreases from the base (left) to the tip (right) of the scale. The hole in the middle corresponds to a region previously milled using the FIB for the thickness measurements.

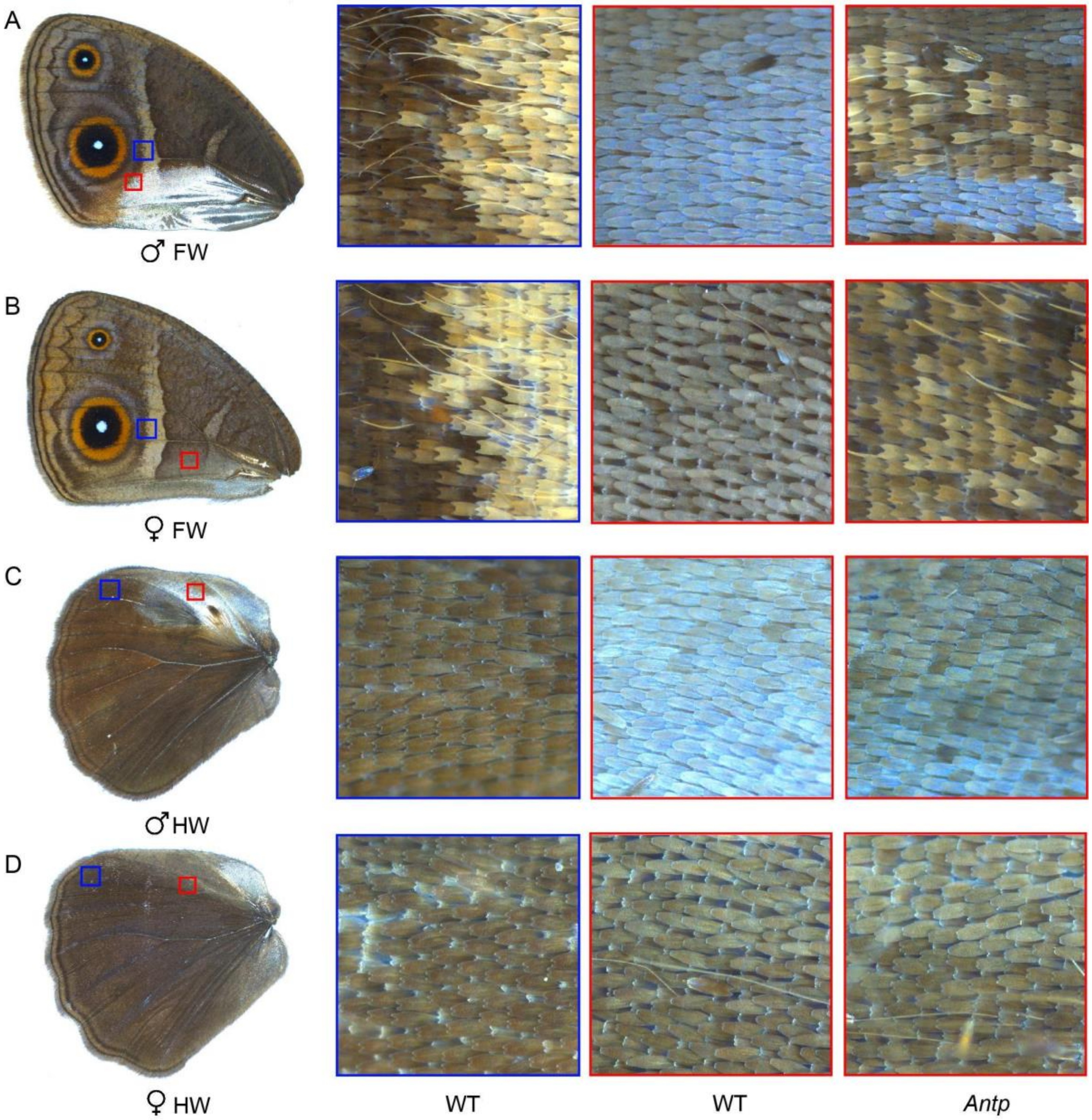


**Supplementary Figure S6: Scale shape in the wildtype and *Antp* crispant wings of male and female *Bicyclus anynana*.** (A,B) Ventral forewings (C,D) Dorsal hindwings. The colored boxes mark the positions on the wing that were imaged. The blue boxes are the control regions where all scales have a dentate distal margin. The red boxes are the regions on the wings that were affected in the *Antp* crispants (dentate scales) as compared to the wildtype (rounded scales).

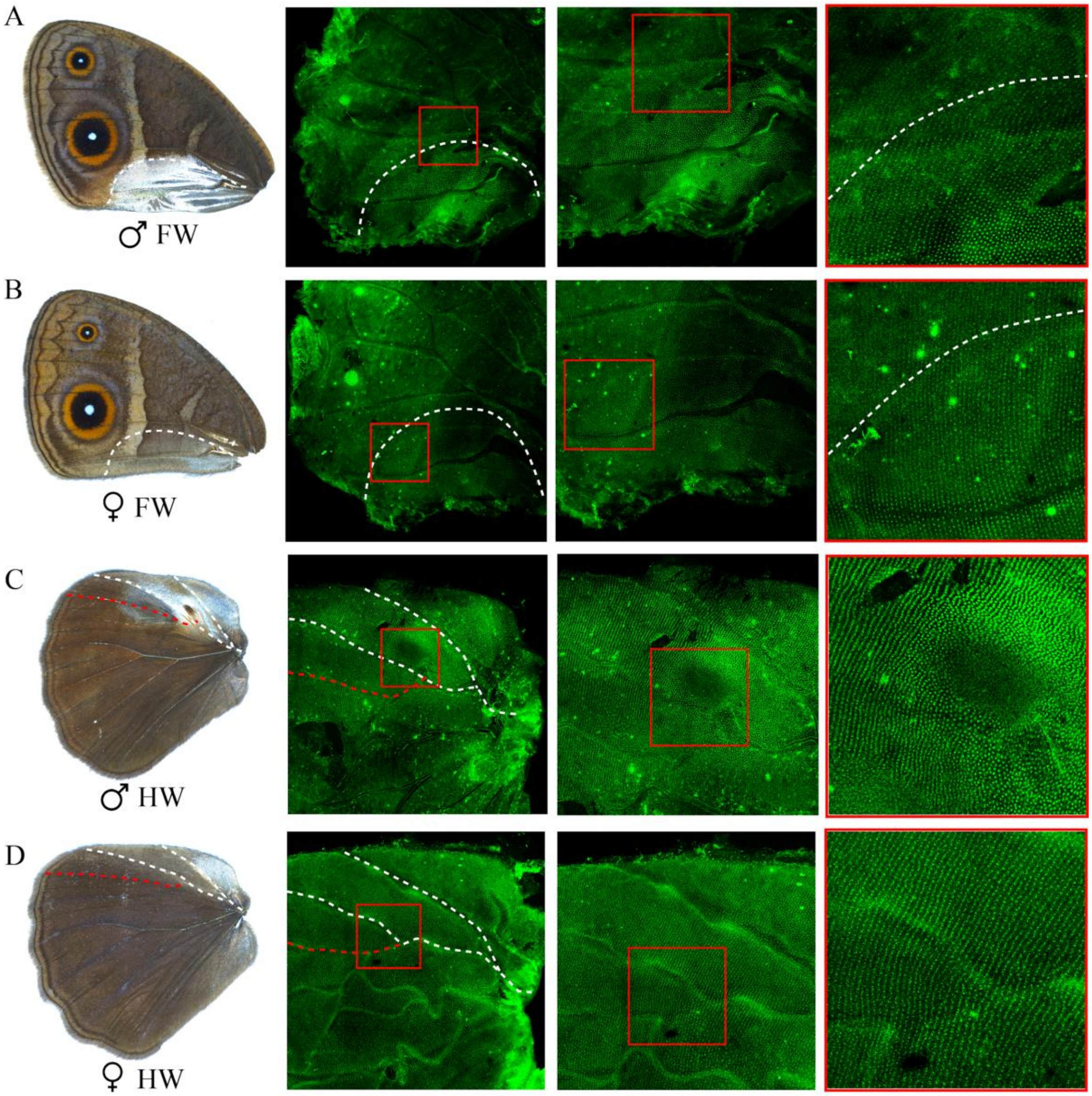


**Supplementary Figure S7: Immunostainings of *Bicyclus anynana* male and female 24-hour pupal wing discs with anti-Antp antibody.** Stainings are shown for (A, B) male and female ventral forewings and (C, D) male and female dorsal hindwings. The white dotted line on the forewings indicates homologous areas on the posterior wings of both sexes where Antp protein is present. The white and red dotted lines on the hindwings highlight homologous veins. The boxed regions are magnified in the last column. Images are the best, illustrative images.

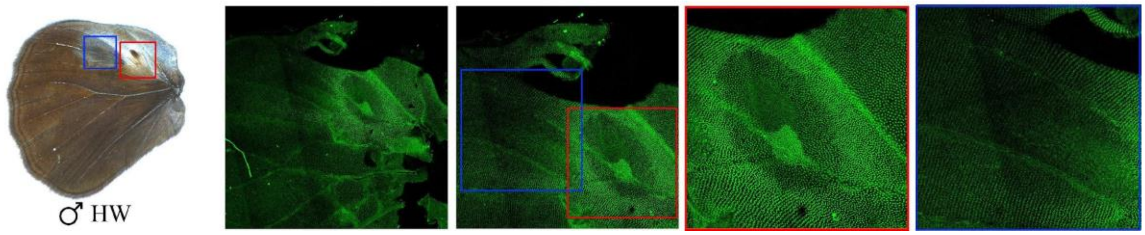


**Supplementary Figure S8: Immunostaining of *Bicyclus anynana* male 48-hour pupal hindwing with anti-Antp antibody.** The red and the blue regions highlight the silver scale region and grey-silver scale region of the hindwings respectively. Antp is present at high levels in the silver scales surrounding the androconia with absence in the brownish-yellow scales in the androconial patch (red box). Low levels of Antp are seen in the grey-silver scale region (blue box).

**Supplementary Table S1: Outcomes of the LME tests comparing area of open upper lamina among wildtype silver scales and between wildtype and crispant scales.**

| Factor | numDF | denDF | F-value | P-value |
|---|---|---|---|---|
| **Fig 1: Area of open upper lamina in wildtype silver scales** | | | | |
| Crispant | 4 | 46 | 17.789998 | **<0.0001** |
| **Fig 2: Area of open upper lamina in *Antp* and *dsx* forewing crispant brown scales compared to wildtype silver scales** | | | | |
| Crispant | 2 | 23 | 709.696 | **<0.0001** |
| **Fig 3: Area of open upper lamina in *apA*, *dsx* and *optix* hindwing crispant brown scales compared to wildtype silver scales** | | | | |
| Crispant | 7 | 82 | 119.0934 | **<0.0001** |
| **Fig 4: Area of open upper lamina in *apA* and *Ubx* ectopic silver scales compared to control brown scales** | | | | |
| Crispant | 4 | 37 | 489.8225 | **<0.0001** |

**Supplementary Table S2: Estimates, 95% CI and adjusted P-values for the multiple comparisons of area of open upper lamina among different scale types obtained from Tukey post-hoc tests.**

| Comparison | Estimate (%) | Std. error | z-value | Pr(>\|z\|) | 95% CI |
|---|---|---|---|---|---|
| **Fig 1: Comparisons of area of open upper lamina among wildtype silver scales** | | | | | |
| FW silver – FW gland == 0 | -0.29793 | 0.09053 | -3.291 | **0.00998** | -0.53360 to -0.06227 |
| HW grey-silver – HW coupling == 0 | 3.07257 | 1.46599 | 2.096 | 0.36091 | -0.74370 to 6.88883 |
| HW silver – HW grey-silver ==0 | -4.69642 | 1.45379 | -3.230 | **0.01236** | -8.48091to -0.91192 |
| **Fig 2: Area of open upper lamina of *Antp* and *dsx* forewing crispant brown scales in comparison to wildtype silver scales** | | | | | |
| WT silver – *Antp* brown == *0* | -41.149 | 1.196 | -34.409 | **<2e-16** | -43.9185 to -38.3797 |
| WT silver – *dsx* brown == 0 | -24.271 | 1.582 | -15.342 | **<2e-16** | -27.9347 to -20.6075 |
| *dsx* brown – *Antp* brown == 0 | -16.878 | 1.983 | -8.511 | **<2e-16** | -21.4706 to -12.2855 |
| **Fig 3: Area of open upper lamina of *apA*, *dsx* and *optix* hindwing crispant brown scales in comparison to wildtype silver scales** | | | | | |
| *optix* brown – WT coupling silver == 0 | 26.7505 | 1.7980 | 14.878 | < **2e-16** | 21.4420 to 32.0591 |
| HW silver region | | | | | |
| WT silver – *apA* brown == 0 | -31.5700 | 2.0911 | -15.097 | < **2e-16** | -37.7439 to -25.3961 |

| | | | | | |
|---|---|---|---|---|---|
| WT silver – *dsx* brown == 0 | -27.8897 | 2.4887 | -11.206 | **< 2e-16** | -35.2376 to -20.5418 |
| HW grey-silver region | | | | | |
| WT grey-silver – *apA* brown == 0 | -25.6673 | 2.9155 | -8.804 | **< 2e-16** | -34.2753 to -17.0593 |
| WT grey-silver – *dsx* brown == 0 | -23.7047 | 2.3649 | -10.023 | **< 2e-16** | -30.6871 to -16.7222 |
| **Fig 4: Area of open upper lamina of *apA* and *Ubx* ectopic silver scales compared to control brown scales** | | | | | |
| *apA* silver – control brown == 0 | -40.7252 | 2.5160 | -16.186 | **< 2e-16** | -47.2815 to -34.1688 |
| *apA* gland – control brown == 0 | -33.4788 | 2.8849 | -11.605 | **< 2e-16** | -40.9965 to -25.9612 |
| *apA* silver – *apA* gland == 0 | -7.2463 | 1.4165 | -5.116 | **3.13e-06** | -10.9376 to -3.5550 |
| *Ubx* silver – control brown == 0 | -39.6885 | 0.9717 | -40.846 | **< 2e-16** | -42.2205 to -37.1564 |

**Supplementary Table S3: Outcomes of the LME tests comparing lamina thicknesses among wildtype silver scales and between wildtype and crispant scales.**

| **Factor** | **numDF** | **denDF** | **F-value** | **P-value** |
|---|---|---|---|---|
| **Fig 1: Lamina thickness of wildtype silver scales** | | | | |
| Crispant | 9 | 537 | 47.8214 | **<0.0001** |
| **Fig 2: Lamina thickness of *Antp* and *dsx* forewing crispant brown scales in comparison to wildtype silver scales** | | | | |
| Crispant | 2 | 225 | 203.8694 | **<0.0001** |
| **Fig 3: Lamina thickness of *apA*, *dsx* and *optix* hindwing crispant brown scales in comparison to wildtype silver scales** | | | | |
| Crispant | 7 | 590 | 87.409 | **<0.0001** |
| **Fig 4: Lamina thickness of *apA* and *Ubx* ectopic silver scales in comparison to control brown scales** | | | | |
| Crispant | 7 | 581 | 201.6783 | **<0.0001** |

**Supplementary Table S4: Estimates, 95% CI and adjusted P-values for the multiple comparisons between lamina thicknesses obtained from Tukey post-hoc tests.**

| **Comparison** | **Estimate (nm)** | **Std. error** | **z-value** | **Pr(>\|z\|)** | **95% CI** |
|---|---|---|---|---|---|
| **Fig 1: Lamina thickness comparisons of wildtype silver scales** | | | | | |

| | | | | | |
|---|---|---|---|---|---|
| FW silver LL – FW silver UL == 0 | 10.5562 | 2.3783 | 4.439 | **0.000408** | 3.1041 to 18.0082 |
| FW gland LL – FW gland UL == 0 | 15.4923 | 2.3472 | 6.600 | **1.85e-09** | 8.1375 to 22.8470 |
| HW silver LL – HW silver UL == 0 | -14.5862 | 3.4778 | -4.194 | **0.001233** | -25.4836 to -3.6889 |
| HW grey LL – HW grey UL == 0 | 6.2368 | 3.4935 | 1.785 | 1.000000 | -4.7097 to 17.1832 |
| HW coupling LL – HW coupling UL == 0 | -22.3861 | 4.7098 | -4.753 | **9.02e-05** | -37.1437 to -7.6284 |
| **Fig 2: Lamina thickness of *Antp* and *dsx* forewing crispant brown scales in comparison to wildtype silver scales** | | | | | |
| *Dsx* brown LL - *Antp* brown LL == 0 | -36.273 | 2.638 | -13.748 | **<2e-16** | -42.4474 to -30.0984 |
| WT FW silver LL – *Antp* brown LL == 0 | -61.299 | 3.113 | -19.693 | **<2e-16** | -68.5842 to -54.0147 |
| WT FW silver LL – *Dsx* brown LL == 0 | -25.027 | 2.899 | -8.633 | **<2e-16** | -31.8105 to -18.2426 |
| **Fig 3: Lamina thickness of *apA*, *dsx* and *optix* hindwing crispant brown scales in comparison to wildtype silver scales** | | | | | |
| *optix* brown LL – WT coupling silver LL == 0 | -7.671 | 4.404 | -1.742 | 1.000000 | -20.9315 to 5.5891 |
| HW silver region | | | | | |
| *apA* brown LL – WT silver LL == 0 | 36.420 | 2.658 | 13.704 | **< 2e-16** | 28.4191 to 44.4213 |
| *dsx* brown LL – WT silver LL == 0 | 33.398 | 3.024 | 11.044 | **< 2e-16** | 24.2934 to 42.5020 |
| HW grey-silver region | | | | | |
| *apA* brown LL – WT grey-silver LL == 0 | 42.917 | 2.917 | 14.715 | **< 2e-16** | 34.1358 to 51.6976 |
| *dsx* brown LL – WT grey-silver LL == 0 | 10.020 | 3.036 | 3.300 | **0.027041** | 0.8794 to 19.1609 |
| **Fig 4: Lamina thickness comparisons of *apA* and *Ubx* ectopic silver scales and control** | | | | | |
| *apA* silver LL – control brown LL == 0 | -59.6307 | 3.1355 | -19.018 | **< 2e-16** | -69.0514 to -50.2099 |

| | | | | | |
|---|---|---|---|---|---|
| *apA* gland LL – control brown LL == 0 | -55.1339 | 3.0188 | -18.264 | **< 2e-16** | -64.2037 to -46.0640 |
| *Ubx* silver LL – control brown LL == 0 | -62.7014 | 2.8596 | -21.926 | **< 2e-16** | -71.2932 to -54.1096 |
| *apA* silver LL – *apA* silver UL == 0 | 7.1981 | 1.9022 | 3.784 | **0.00432** | 1.4831 to 12.9132 |
| *apA* gland LL – *apA* gland UL == 0 | 10.9035 | 1.6804 | 6.489 | **2.42e-09** | 5.8549 to 15.9522 |
| *Ubx* silver LL – *Ubx* silver UL == 0 | 15.6452 | 2.7357 | 5.719 | **3.00e-07** | 7.4259 to 23.8645 |
